# Transcription-replication interactions reveal principles of bacterial genome regulation

**DOI:** 10.1101/2022.10.22.513359

**Authors:** Andrew W. Pountain, Peien Jiang, Tianyou Yao, Ehsan Homaee, Yichao Guan, Magdalena Podkowik, Bo Shopsin, Victor J. Torres, Ido Golding, Itai Yanai

## Abstract

Organisms determine the transcription rates of thousands of genes through a few modes of regulation that recur across the genome. These modes interact with a changing cellular environment to yield highly dynamic expression patterns. In bacteria, the relationship between a gene’s regulatory architecture and its expression is well understood for individual model gene circuits. However, a broader perspective of these dynamics at the genome-scale is lacking, in part because bacterial transcriptomics have hitherto captured only a static snapshot of expression averaged across millions of cells. As a result, the full diversity of gene expression dynamics and their relation to regulatory architecture remains unknown. Here we present a novel genome-wide classification of regulatory modes based on each gene’s transcriptional response to its own replication, which we term the Transcription-Replication Interaction Profile (TRIP). We found that the response to the universal perturbation of chromosomal replication integrates biological regulatory factors with biophysical molecular events on the chromosome to reveal a gene’s local regulatory context. While the TRIPs of many genes conform to a gene dosage-dependent pattern, others diverge in distinct ways, including altered timing or amplitude of expression, and this is shaped by factors such as intra-operon position, repression state, or presence on mobile genetic elements. Our transcriptome analysis also simultaneously captures global properties, such as the rates of replication and transcription, as well as the nestedness of replication patterns. This work challenges previous notions of the drivers of expression heterogeneity within a population of cells, and unearths a previously unseen world of gene transcription dynamics.

## Introduction

Our ability to understand and manipulate bacteria, from design of synthetic regulatory circuits^1^ to determining how bacterial pathogens establish and maintain infection in their hosts, demands a sophisticated understanding of gene regulatory processes. Bacterial gene regulation occurs primarily at the level of transcription^2^, but while decades of research has produced a wealth of knowledge about RNA polymerase and its interactions with promoters, repressors, and activators of transcription, this work is primarily based on measurements averaged across a population of millions of cells. Therefore, much is still unclear about how transcription takes place in individual cells in the context of a constantly changing cellular environment^3^. In rapidly proliferating cells, transcription occurs on a chromosome that is under continuous replication^4,5^. However, although there has been some exploration of the effects of replication on individual genes^6,7^, the transcriptome-wide consequences of this perturbation are unknown^8,9^. Measuring global gene expression during the replication cycle has traditionally been hampered by the requirement for analysis of synchronized populations at a bulk level, limiting this analysis to organisms such as *Caulobacter crescentus*^10–12^ where natural biological features facilitate synchronization, or to populations synchronized by batch synchronization methods such as starvation^13^ or temperature shift^14^ that may be both of questionable efficacy and liable to introduce artefacts^15^.

Here we combined state-of-the-art bacterial single cell RNA sequencing (scRNA-seq)^16–19^ with a new cell cycle analysis framework to reveal extensive transcriptional variation during the cell cycle in two unrelated species – the model organism and Gram-negative rod *Eschericha coli* (*E. coli*), and the Gram-positive coccus *Staphylococcus aureus* (*S. aureus*), both major bacterial pathogens. We identified first a global replication-dependent pattern that depends on a gene’s chromosomal location, then developed a predictive computational analysis framework to reveal diverse types of divergence from this pattern. In *E. coli*, we found an effect of a gene’s position within its operon on expression dynamics that is largely absent in *S. aureus*. Other genes diverged from the expected pattern in both amplitude and timing of their expression in ways that are sensitive to gene-specific factors such as repression state. Therefore, while DNA replication introduces a universal perturbation, how individual genes respond to this perturbation depends on their local regulatory context, providing a new lens through which to understand the behavior of genes at their native loci.

### Global gene expression in proliferating bacterial populations is shaped by chromosomal organization

To investigate transcriptional heterogeneity in proliferating bacterial populations, we applied a recently-described scRNA-seq method, PETRI-seq^16^, to 73,053 individual *S. aureus* cells in exponential phase (Fig. 1A). *S. aureus* is an important human pathogen, yet little is known about heterogeneous gene expression dynamics within its populations. We detected on average 135 transcripts per cell (Fig. S1A), an increase on the 43 transcripts per cell previously published for this species with this method^16^. As the data are very sparse, we denoised them using the single-cell variational inference (scVI) method, an unsupervised deep learning approach^20^. Studying gene-gene correlations, we recovered the expected covariance of genes within operons (Fig. 1B). However, when we investigated gene-gene correlations on a genomic scale, we discovered a striking ‘X-shaped’ pattern of gene expression covariance (Fig. 1C, Fig. S2A). The central ‘X’ of this pattern reflects symmetry around the origin of replication, meaning that genes equidistant from the origin on each side of the chromosome correlate with each other. Beyond the ‘X’ itself, however, we observed an additional correlation directly between genes at the origin and terminus (Fig. 1C). This pattern was strengthened by averaging expression into 50 kb bins by chromosome position (Fig. 1C), and was reproducible in a second independent dataset under the same conditions of 21,257 cells (Fig. S2C). It was detectable even without the use of scVI, although the signal was noisier (Fig. S2B). The pattern was abolished when we studied 55,894 cells in stationary phase, suggesting that it is a property of proliferating cells (Fig. 1D).

**Figure 1:**
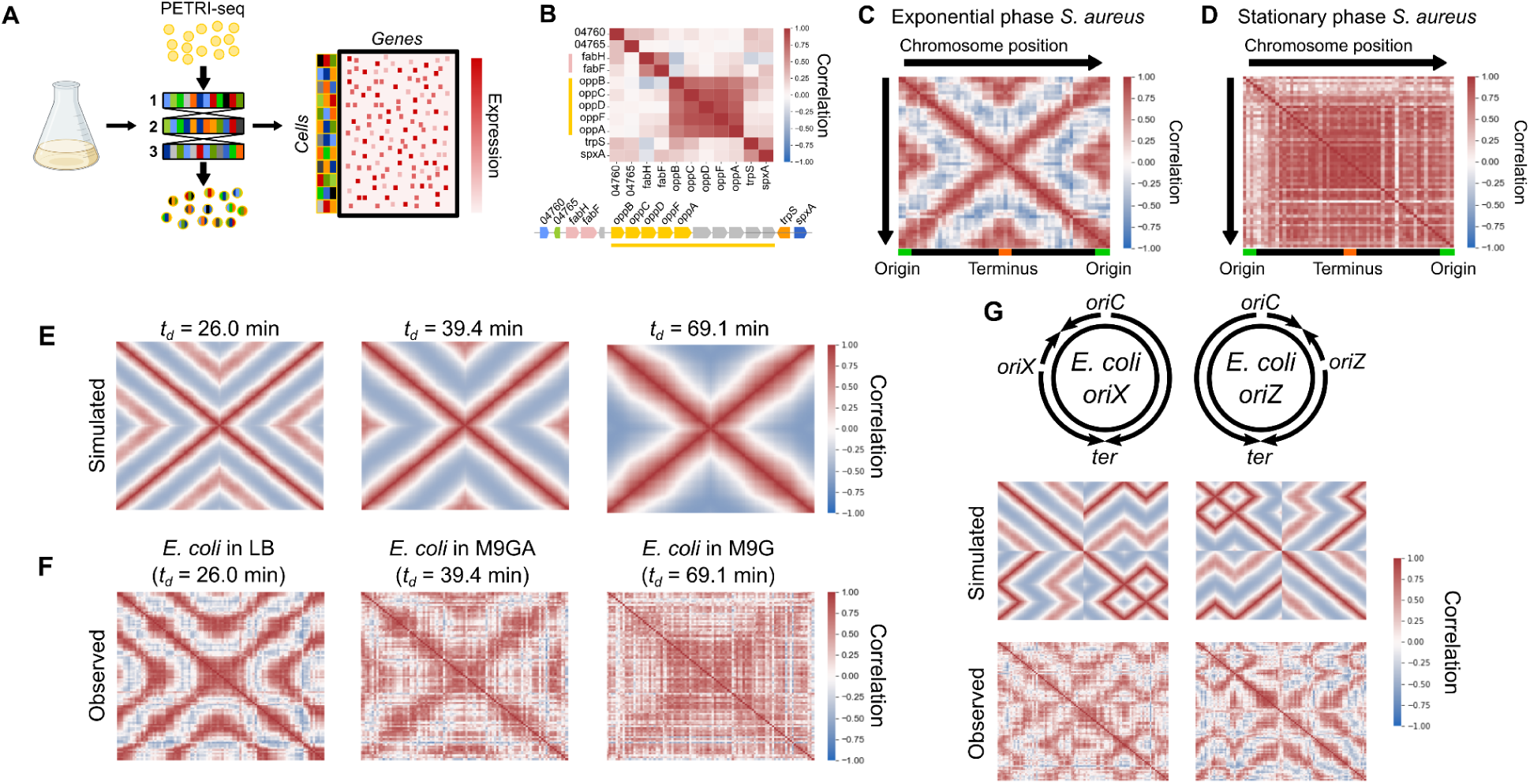
scRNA-seq reveals a global pattern of replication-associated gene covariance. **A)** PETRI-seq workflow^16^. Bacterial cells were fixed and permeabilized, then subjected to three rounds of cDNA barcoding to give transcripts of each cell a unique barcode combination. This method is highly scalable to multiple samples and tens of thousands of cells. **B)** Local operon structure is captured by gene-gene correlations (Spearman’s *r*). Operons are indicated by shared colors of genes. Gray genes indicate those removed by low-count filtering. Names of *SAUSA300_RS04760* and *SAUSA300_RS04765* are truncated. **C & D)** Global gene-gene correlations reflect chromosomal position in **(C)** exponential phase and **(D)** stationary phase *S. aureus*. Spearman correlations were calculated based on scVI-smoothed expression averaged in 50 kb bins by chromosome position. **E)** Simulated correlation patterns in unsynchronized *E. coli* populations at three different growth rates. **F)** Spearman correlations between scaled data averaged into 50 kb bins, as for **(C)** but for *E. coli* grown at three growth rates. **G)** Introducing ectopic origins of replication in *E. coli* leads to predictable perturbations in gene expression heterogeneity. *Top*: schematic of predicted replication patterns based on previous studies^21–23^. *Middle*: Predicted correlation patterns based on the copy number simulation. *Bottom*: Real correlation patterns in *oriX* and *oriZ* mutant strains, as in **(C)**. Heatmaps of correlations without chromosome position-dependent binning are shown in Fig. S2D.

As we observed correlations among genes that are equidistant from the origin of replication and cells in stationary phase did not show such correlations, we hypothesized that the ‘X-shaped’ pattern reflects the effect of DNA replication on gene expression. In the model organism *E. coli*, replication patterns are growth rate-dependent: at high rates of proliferation, overlapping cycles of replication occur simultaneously, whereas at slower proliferation rates one round of replication is completed before the next one begins^4,24^. This arises because the ‘C-period’, the time for one complete round of replication from the origin to the terminus, remains approximately constant and can be greater than the doubling time^4,24^. The effect of replication on gene expression covariance should reflect this. To test this, we therefore measured the doubling times (*t_d_*) of *E. coli* grown at 37 °C in three medium conditions (Fig. S3A): LB (26.0 ± 1.3 min), M9 minimal medium with glucose and amino acids (M9GA, 39.4 ± 2.3 min), and M9 medium with glucose only (M9G, 69.1 ± 9.8 min). We next developed a simulation to predict correlation patterns arising from gene dosage in cells proliferating with these doubling times (Fig. 1E & Fig. S4). At an intermediate growth rate (*t_d_* = 39.4 min), we predicted a correlation pattern similar to that observed for *S. aureus* (Fig. 1C). However, simulating faster growth produced a nested “multi-X” pattern resulting from overlapping cycles of replication, and slower growth greatly reduced origin-terminus correlations (Fig. 1E).

When we compared these predictions to the observed data for *E. coli* grown under the three conditions, we observed a close correspondence between simulated and observed expression patterns (Fig. 1F). Correlations became less defined at slower growth rates, although this may reflect technical noise due to lower transcript counts (Fig. S1B), resulting from lower RNA content at slower growth rates^25^. The correlation pattern of *E. coli* grown in M9G, the slow-growth condition, further resembled bulk RNA-seq of synchronized *C. crescentus* (Fig. S4C)^11^, a species that undergoes a single round of replication prior to asymmetric division^10^, which is a similar situation to that of slower-growing *E. coli*. Next, we reasoned that if this pattern is driven by the effect of gene copy number on expression levels (as assumed in our simulation), we also expect to find a relationship between origin distance and expression levels. Indeed, despite high variation in intrinsic promoter activity, we found that on average gene expression decreased with distance from the origin, and this effect was stronger at faster growth rates^26^ (Fig. S5). Finally, while these patterns could theoretically arise due to reads from contaminating genomic DNA, multiple lines of evidence from the data (Fig. S6), as well as our observation of the X-shaped pattern in a previously published dataset of bulk RNA from synchronized *C. crescentus*^11^ (Fig. S4C), demonstrate that this is very unlikely to be the case and support our interpretation that the observed patterns are driven by the effect of DNA replication on mRNA abundance.

To further test our ability to predict global correlations from expected replication patterns, we examined strains in which normal replication is perturbed. We compared wild-type *E. coli* grown in LB to two strains with ectopic origins of replication at either 9 o’clock (*oriX*) or 3 o’clock (*oriZ*) positions in addition to *oriC*^21–23^. In these strains, replication initiates simultaneously at both native and inserted origins, while ending at the same terminus, *ter*^21^. Our simulation predicted perturbed correlation patterns that were almost mirror images of each other, given that the ectopic origins of the mutants we chose were nearly equidistant from *oriC* on each side of the chromosome (Fig. 2G). Again, we found that the observed patterns matched closely with our predictions (Fig. 2G). These results support the notion that DNA replication kinetics produce a predictable effect on transcriptional heterogeneity within a population of proliferating bacteria, and that this effect is sensitive to growth rate and genetic perturbations.

**Figure 2:**
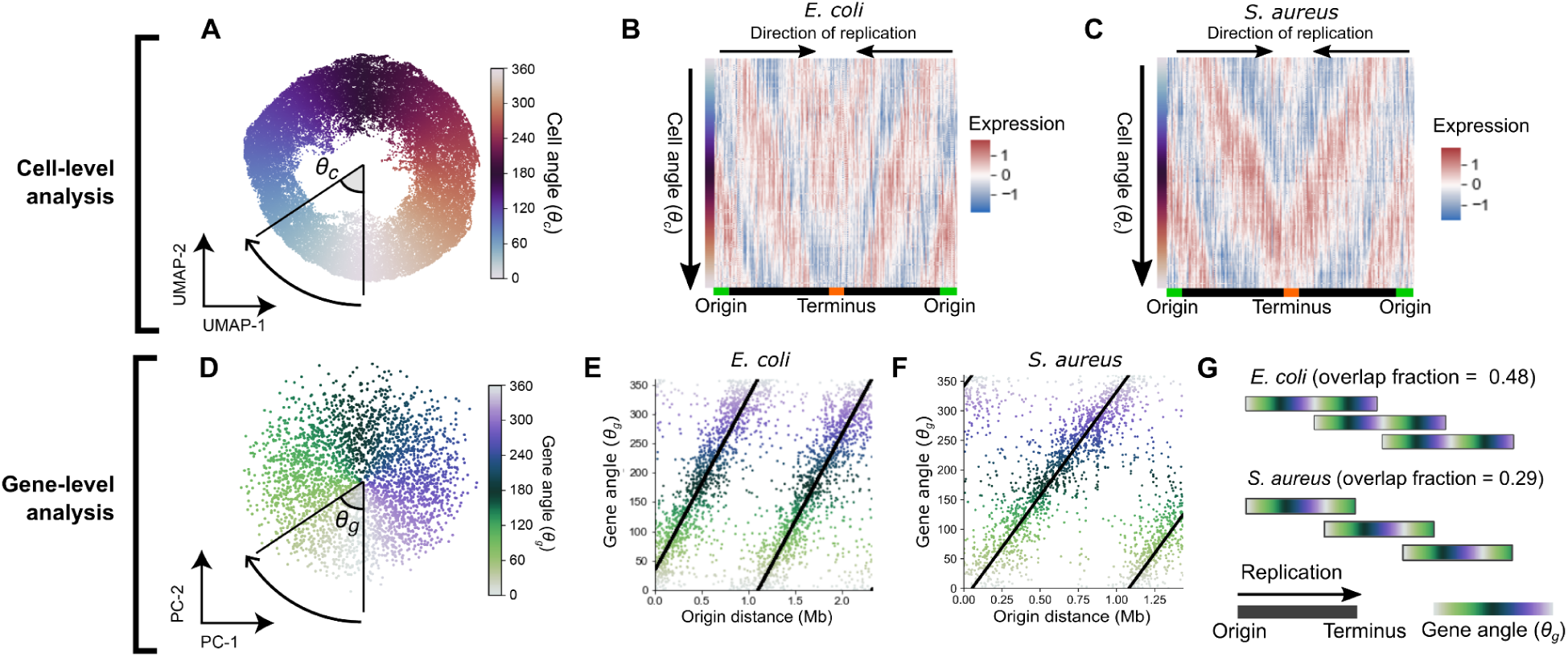
Ordering expression by cell angle and gene angle provides a quantitative description of cell cycle gene expression. **A)** UMAP of LB-grown *E. coli* with expression averaged in 100 kb bins by chromosome position. Cell angle *θ_c_*is the angle between UMAP dimensions relative to the center. For UMAP without averaging, see Fig. S7A. **B & C)** Heatmap of scaled gene expression in *E. coli* **(B)** or *S. aureus* **(C)** averaged in 100 bins by *θ_c_*. **D)** Derivation of gene angle *θ_g_* in LB-grown *E. coli*. Principal component analysis was performed on the transpose of the matrix in **(B)**, and *θ_g_* was defined as the angle between principal components (PCs) 1 and 2. Genes form a wheel in UMAP (Fig. S7C). **E & F)** The relationship between *θ_g_* and origin distance for *E. coli* grown in LB **(E)** and *S. aureus* grown in TSB **(F)**. **G)** Predicted replication patterns in LB-grown *E. coli* (*t_d_* = 26.0 ± 1.3 min) and *S. aureus* (*t_d_* = 24.9 ± 0.6 min). Overlapping rounds of replication lead to shared *θ_g_* in simultaneously-replicated chromosomal regions. Note that greater overlap in replication rounds is observed for *E. coli* than for *S. aureus*.

### The effect of chromosomal replication on transcription facilitates resolution of bacterial gene expression by cellular replication state

Since DNA replication exerts a strong influence over gene expression, we reasoned that this effect can be used to resolve a cell’s position within the replication cycle given only its transcriptome. To examine the distribution of cellular states in a population of cells, we projected gene expression measurements of LB-grown *E. coli* cells in two dimensions by uniform manifold approximation and projection (UMAP^27^). Cells arranged into a “wheel” shape (Fig. 2A) when we performed UMAP on expression averaged by chromosomal position (which was found to strengthen global correlation patterns, Fig. 1C). To determine the order of cells along this wheel, we calculated cells’ angle *θ_c_*between UMAP coordinates (Fig. 2A). Examining gene expression as a function of *θ_c_*, we observed waves of gene expression progressing from the origin to the terminus (Fig. 2B), suggesting that cells’ positions on this wheel reveal their replication state. Performing equivalent analysis to resolve replication states in *S. aureus*, we observed a similar pattern (Fig. 2C, Fig. S7B). These data suggest that we can infer a cell’s replication state from the transcriptome alone, and that this holds across different bacterial species.

As we observed that the expression of most genes is strongly influenced by a cell’s replication state, we reasoned that we should also be able to order genes by their timing of expression within the cell cycle and that this would generally reflect their order of replication. To do this, we projected the genes themselves into two dimensions to derive a gene angle, *θ_g_* (Fig. 2D). We observed a close relationship between the order of genes by *θ_g_* and the distance from the origin of replication in both *E. coli* and *S. aureus* (Fig. 2E & F), suggesting that *θ_g_* does indeed capture the order of replication. However, we also observed that the period of *θ_g_* (i.e. the chromosomal distance associated with a 360° rotation) was less than the full origin-terminus distance, meaning that genes at multiple positions on the origin-terminus axis had the same *θ_g_* value. We can interpret this to mean that at high growth rates, overlapping rounds of replication lead to simultaneous replication of genes at multiple distances from the origin. Furthermore, we observed that in *E. coli*, the gradient of change of *θ_g_*with respect to origin distance decreased with slowing growth rate (Fig. S7D & F). We can use this gradient to infer two parameters about the replication pattern. Firstly, this gradient provides an estimate of the average DNA polymerase speed. For *E. coli* in LB, this estimate was 780 bp/s (Fig. S7F), very close to previously reported values of ∼800 bp/s^28,29^. Secondly, the gradient can also be used to estimate an “overlap fraction” (Fig. 2G), the fraction of one round of replication happening before the previous one has finished. When we compared *E. coli* at different growth rates, we observed that, in line with expectations^4,24^, decreasing proliferation speed in *E. coli* is associated with reduced overlap in rounds of replication (Fig. S7E), while the average DNA polymerase speed (and hence the C-period) remains roughly consistent (Fig. 7F). In *S. aureus*, the reduced size of its genome (2.9 Mb vs 4.6 Mb in *E. coli*) explains why, despite similar proliferation rates and DNA polymerase speeds (Fig. S7F), less overlap in rounds of replication is observed than *E. coli* (Fig. 2G). Therefore, the gene angle *θ_g_* and its relationship to distance from the replication origin provide a quantitative and interpretable description of the relationship between gene expression and global replication patterns.

Finally, the two parameters we introduce here – the cell angle *θ_c_* and the gene angle *θ_g_*(Fig. S7 G & H) – led us to construct an inference model to predict the expression of a given gene (by *θ_g_*) at a given point in the cell cycle (by *θ_c_*), based on global replication-dependent trends (Fig. S8). Thus based on a given pattern of gene expression, the model infers the state of the cell along the cell cycle; conversely, for a particular cell cycle state, the model infers an expected gene expression pattern based solely on a gene’s distance from the origin (and hence replication timing). Overall, we found a moderate correlation of this prediction with the observed data (Pearson’s *r* = 0.59, Fig. S9A), and subtraction of this prediction from the observed data eliminated the global correlation pattern (Fig. S9B), confirming that our model effectively captured position-dependent gene expression trends.

### The global consensus pattern of gene expression reflects a replication-dependent gene dosage effect

We next sought to confirm that the transcriptional dynamics we inferred from the scRNA-seq data represent cell cycle-dependent gene expression. To do this, we first identified three operons whose genes’ expression closely fits the model-predicted pattern (Fig. 3A), then compared our measurements for genes within the selected operons to cell cycle-dependent gene expression measurements obtained using single molecule fluorescence *in situ* hybridization (smFISH)^6,30^. Overall, population-averaged expression measurements from the two methods were in close quantitative agreement (Fig. S10D). The smFISH approach resolves cell cycle by using cell length to infer cell age, thus defining the cell cycle relative to *division* timing^6^. By contrast, we defined cell angle *θ_c_*= 0 to be the assumed time of *replication initiation* (see Materials & Methods). As expected given these differing “start” points, we observed a phase shift in expression profiles between the two methods that was consistent across genes (Fig. S10E). Modeling of total DNA content as a function of cell length supported that this phase shift was roughly consistent with our choice of *θ_c_* = 0 as the point of replication initiation (Fig. S10F), albeit with some discrepancy (see Materials & Methods).

**Figure 3:**
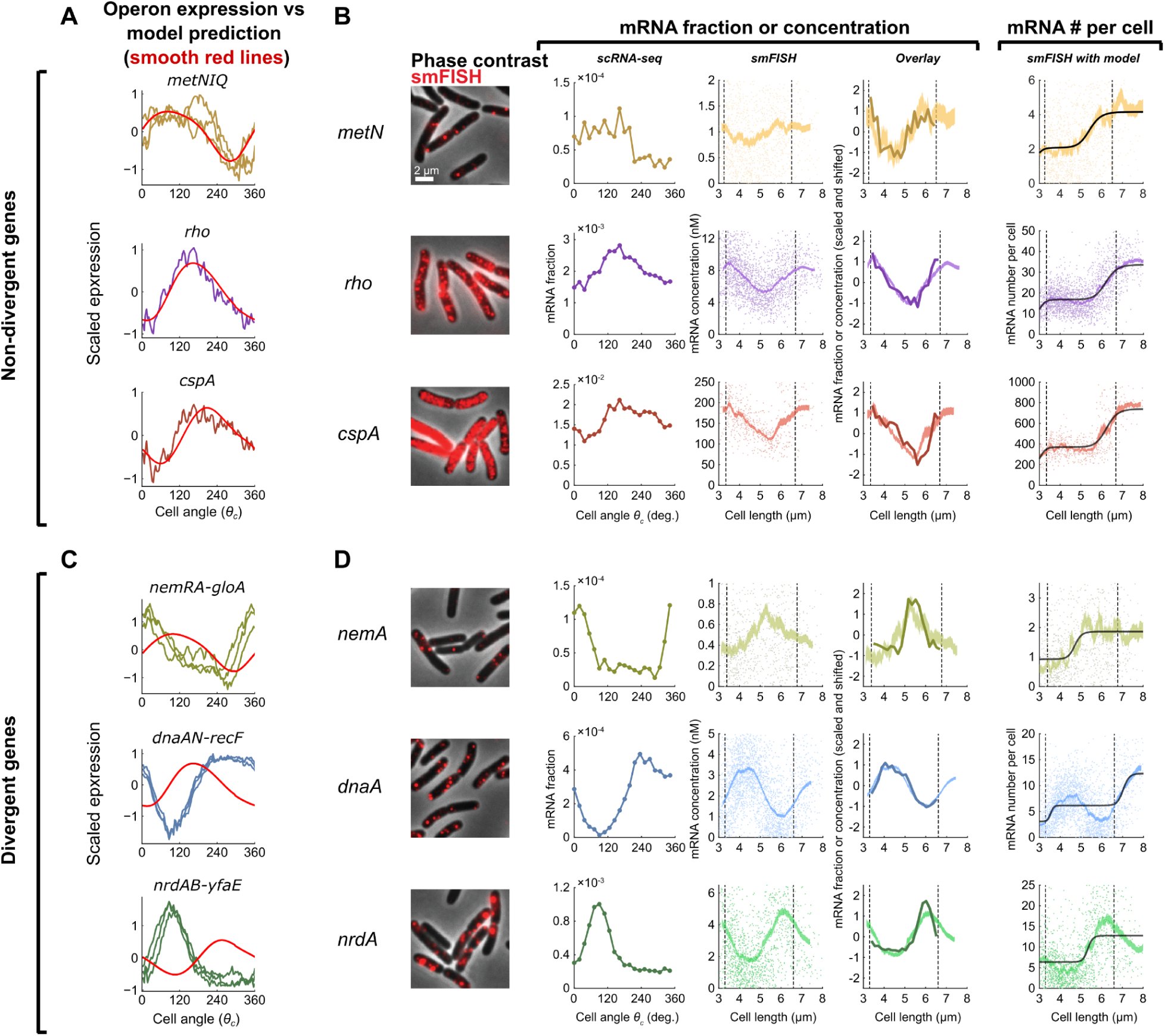
Genes show a spectrum of divergence from a dosage-driven consensus pattern. **A)** Expression of genes in operons that conform to the consensus pattern across 100 bins averaged by *θ_c_*. Expression is *z*-scores derived from scVI (jagged lines) or predicted as a replication effect (smooth, red lines). **B)** Comparison of scRNA-seq and smFISH data for genes within non-divergent operons. From left to right: *1)* Microscopy images of *E. coli* cells labeled using smFISH against the indicated gene (*cspA* is visualized with alternative contrast; for negative control see Fig. S10A); *2)* scRNA-seq expression shown as fraction of total cellular mRNA (expression is averaged in 100 bins by *θ_c_*); *3)* mRNA concentration, measured using smFISH, as a function of cell length. Single-cell data (scatter plot) was binned by cell length (shaded curve, moving average ± SEM, 10% sample size per bin). Dashed lines indicate the twofold length range where most cells reside, used to infer the mean values at birth and division; *4)* Alignment of scaled data from smFISH and scRNA-seq measurements; *5)* Absolute mRNA copy number, measured using smFISH, as a function of cell length. Single-cell data was processed as in column 3 (5% sample size per bin). Black line, fit to a sum of two Hill functions, corresponding to two gene replication rounds. **C)** Expression of divergent genes compared to model predictions (as in **(A)**). **D)** Comparison of scRNA-seq and smFISH as in **(B)** but for divergent genes. See Material and Methods for further details.

By correcting for this phase shift between methods, we aligned the scRNA-seq profile to that of the smFISH data (Fig. 3B). In doing so, we observed that expression dynamics inferred by the two methods were highly correlated, confirming that our scRNA-seq approach captures cell cycle-dependent expression. Moreover, while our scRNA-seq measurements capture only relative expression of a gene among total cellular mRNA, our smFISH experiments additionally provide us absolute abundance. This revealed a discrete twofold stepwise increase in expression (Fig. 3B), consistent with genes that are sensitive to gene dosage but otherwise exhibit constant expression^6^. These observations support an interpretation that the model-predicted pattern corresponds to cell cycle expression variation driven by gene dosage.

### Genes that diverge from the global consensus pattern exhibit gene dosage-independent features

While many genes conform to this gene dosage-driven expression pattern, others differ from it in a variety of ways. To identify genes that diverged from the expected pattern, we used the predictive model developed above to derive a score for divergence, which we found to be correlated between replicates for genes that showed high variance across the cell cycle (Pearson’s *r* = 0.80, Fig. S9D). We then focused on three operons whose genes strongly diverged from the expected pattern, two of which were involved in replication initiation and elongation (*dnaAN-recF* and *nrdAB-yfaE*, respectively) and one involved in the response to reactive electrophilic species (*nemRA-gloA*)^31–33^. Divergent genes within the same operon showed highly similar expression profiles (Fig. 3A & C), but showed reproducible patterns that differed markedly from predictions (Fig. 3C), while also closely aligning with smFISH measurements (Fig. 3D, Fig. S11). Moreover, both scRNA-seq and smFISH showed that the amplitude of cell cycle expression (i.e. the relative change between cell cycle minimum and maximum expression) was higher for these divergent genes than the non-divergent ones (Fig. S10G). Finally, absolute mRNA copy number measurement demonstrated that unlike the non-divergent genes, *dnaA* and *nrdA* do not conform to a dosage-related step function (Fig. 3D). Taken together, therefore, we observe that genes diverging from the predicted global pattern do so in both shape and timing of expression profile, as well as amplitude, suggesting that additional factors beyond gene dosage drive their expression dynamics. This motivated us to investigate further the factors shaping the divergences in each species.

### The location of genes within operons influences cell cycle expression dynamics in *E. coli*

We first sought to determine what contributes to differential timing of expression profiles among divergent genes. In *E. coli*, we observed the systematic bias that the majority of divergent genes showed delayed expression dynamics relative to predictions (*θ_g_* is more “clockwise” than *θ_g-pred_*, Fig. 4A). Many of these genes were encoded in large operons, such as those involved in energy biogenesis (e.g. *nuo* and *atp* operons) and cell surface synthesis (e.g. the *mraZ-ftsZ* operon). We found that genes with a more distal position within these operons exhibited a greater delay (Fig. 4B, Fig. S13A). Moreover, this delay was relative to the timing of replication: in genes whose replication-predicted pattern changed in the *oriZ* mutant, expression shifted in this strain so that the delay was relative to this new replication time (Fig. 4B). Across all genes, we observed a modest but highly significant correlation between this “angle difference” and distance from the transcriptional start site (TSS) (Fig. 4C). We hypothesized that this delayed phenotype arises due to the time for RNA polymerase (RNAP) to reach genes after replication by DNA polymerase (DNAP) has occurred. The speed of RNAP has previously been estimated as 40 nt/s^6,34^, much slower than the ∼800 nt/s speed for DNAP (^28,29^ and Fig. S7F). By performing linear regression to measure the angle difference/transcriptional distance relationship (Fig. 4C) and converting *θ_g_* into time by assuming that 360° is equivalent to one doubling time of 26 min, we infer that distance from the TSS is associated with a delay that is consistent an with average RNAP speed of 32 nt/s (38 nt/s in a second replicate, Fig. 13C). Therefore, our data support the hypothesis that when a gene is replicated, the time for expression to increase to the higher-expressed state (due to higher gene dosage) correlates with the time for RNAP to reach that same gene after transcription from the replicated locus restarts.

**Figure 4:**
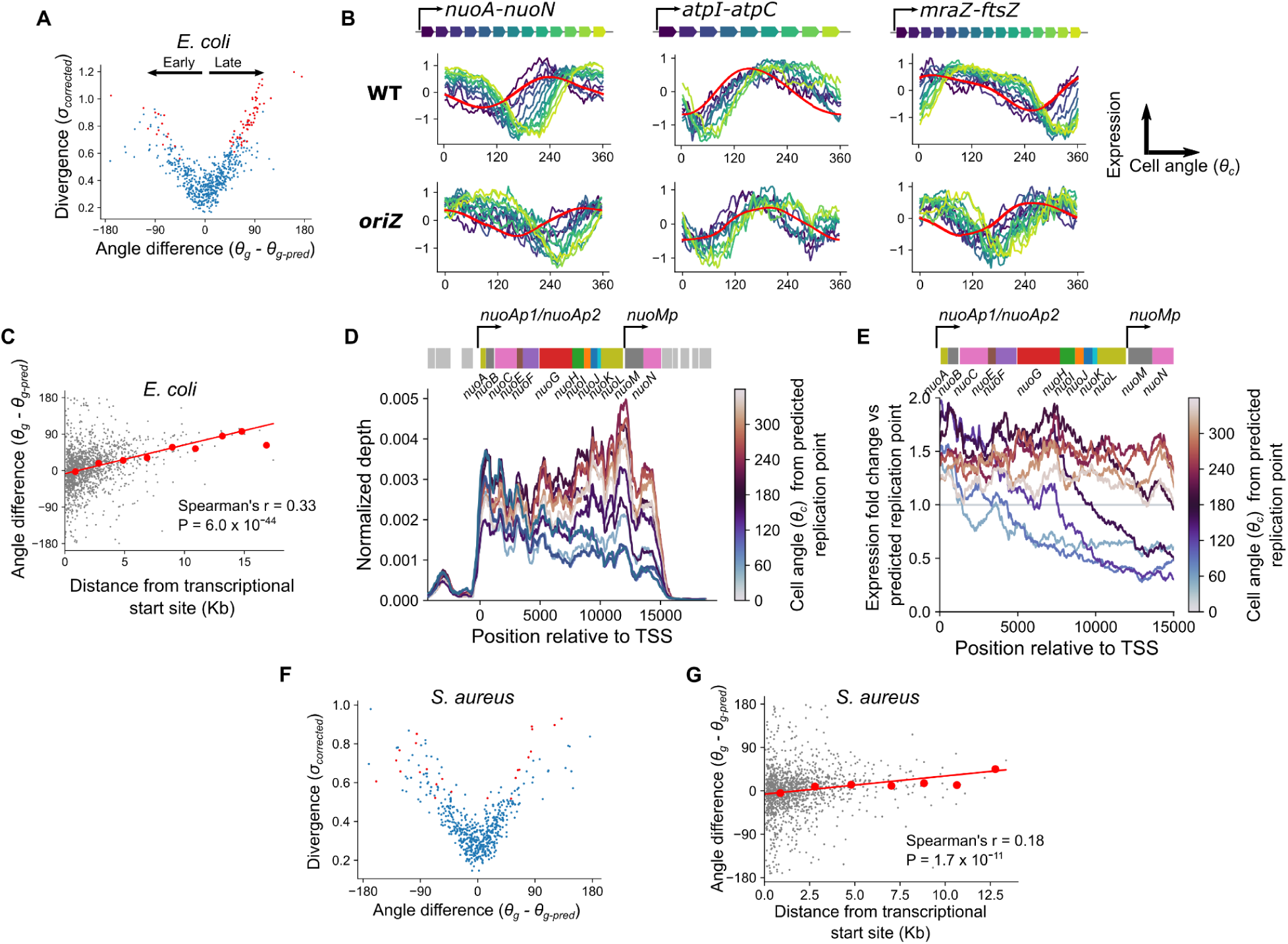
A gene’s position within its operon produces a characteristic delay in expression dynamics in *E. coli* but not *S. aureus*. **A)** Plot of divergence from predictions against the difference between predicted and observed angles in *E. coli*, with divergent genes in red. Angle difference therefore represents whether a gene is expressed earlier or later than expected, as indicated by the black arrows. **B)** Cell cycle expression plots for operons showing “delayed” genes as in Fig. 3A & C but colored by position within the operon. Model-predicted expression is represented in red. Shown for WT and the *oriZ* mutant. **C)** Plot of maximum distance from a transcriptional start site against difference between predicted and observed angles in *E. coli*. Red line indicates the linear model fit and red points indicate averages of 2 kb bins. **D)** Normalized per-base read depth at the *nuo* operon locus for cells averaged in 10 bins by cell angle, *θ_c_*. Traces are smoothed by a 1 kb centered rolling mean and colored by mean cell angle relative to the predicted timing of gene replication (see Materials & Methods). The *nuo* operon structure is indicated by the schematic above, with the surrounding genes in grey. **E)** Per-base read depth as shown in **(D)** for the *nuo* operon, but with expression shown as fold-change relative to expression at the predicted time of gene replication. **F)** Plot of divergence from predictions against the difference between predicted and observed angles, as in **(A)** but for *S. aureus*. **G)** Plot of maximum distance from a transcriptional start site against difference between predicted and observed angles, as in **(B)** but for *S. aureus*.

To further understand the nature of this transcriptional distance effect, we focused on a single operon encoding the NADH dehydrogenase I complex (*nuo*). We observed a delayed effect that increased with distance from the major TSS for this operon, similar to the delay recently observed for this operon in response to transcription initiation inhibition by rifampicin^2^ (Fig. 4B). Additionally, however, where coverage of genes close to the TSS increase in expression immediately after the predicted time of gene replication, coverage at the distal end of the operon dropped sharply before recovering to a higher level (Fig. 4D & E). A similar drop was observable for genes far from the TSS in the *mraZ-ftsZ* operon (Fig. S13B). A potential mechanistic explanation for this is as follows: since passage of the replication fork leads to local disruption of ongoing transcription^35^, genes at the distal end of a transcript are more likely to experience disruption before their transcription can be completed, and there will be a longer delay before new transcription of these genes resumes after replication. This in turn would lead to a post-replication *drop* in expression of genes far from the TSS, compared to an immediate rise in genes close to it. In turn, this would lead to higher amplitude of expression (maximum vs minimum expression) within the cell cycle for genes far from their TSS. Consistent with this, we observed a weak but significant correlation in *E. coli* between genes’ distance from their TSS and their amplitude of expression (Spearman’s *r* = 0.16, P = 2.3 x 10^-10^) (Fig. S13D). We note that many long operons in *E. coli* (e.g. the *nuo* and *mraZ-ftsZ* operons described here, Fig. 4D, Fig. S13B, and ^36^) contain internal promoters, and we suggest that these may contribute to expression by buffering the effects of replication-associated abortive transcription in long operons.

Finally, we asked whether similar trends could be observed in *S. aureus*. In contrast to *E. coli*, we did not observe an excess of “delayed” genes among the divergent genes (Fig. 4F). Moreover, the relationship between operon position and the difference between observed and predicted gene angles was weaker in this species (Fig. 4G), with no observable effect of distance from the TSS on expression amplitude (Spearman’s *r* = 0.01, P = 0.73) (Fig. S13D). From the gradient of this relationship, we predicted that distance from the TSS introduces a delay of 64 nt/s (92 nt/s and 59 nt/s in additional replicates, Fig. S13C). These differences between species persisted even when operons were redefined according to simpler criteria (tandemly arrayed genes with intergenic distance less than 40 bp^37^, Fig. S13E). One potential explanation for this is that if the RNAP processivity rate were faster in *S. aureus* than in *E. coli*, the delay before it reached genes at the distal end of operons would be far less pronounced. In keeping with this, experimental measurement of RNAP by a reporter system in *Bacillus subtilis*, like *S. aureus* a firmicute of the order Bacillales, suggested that it was substantially faster (75–80 nt/s) than its counterpart in *E. coli* measured by the same method (∼48 nt/s)^38,39^. Therefore, the interplay between DNAP and RNAP processivity may lead to species-specific effects of operon position on cell cycle expression dynamics.

### Repressed genes exhibit higher amplitude pulses in cell cycle gene expression

Although the position of genes within operons explains the delayed expression pattern observed in *E. coli*, it can not explain divergent patterns for many other genes in both *E. coli* and *S. aureus*. Therefore, we investigated more closely the shape of cell cycle expression curves for those genes that had reproducible dynamics across replicates (Fig. S14B). To compare genes at different chromosomal loci, we introduced an alignment procedure whereby time is represented as progression by cell angle relative to a gene’s predicted replication time, *θ_c-rep_* (Fig. 5A). Most genes rise rapidly (presumably due to a doubling of gene dosage) before declining as a relative fraction of the transcriptome. Many genes, however, exhibited patterns that could not be explained by gene dosage effects alone.

**Figure 5:**
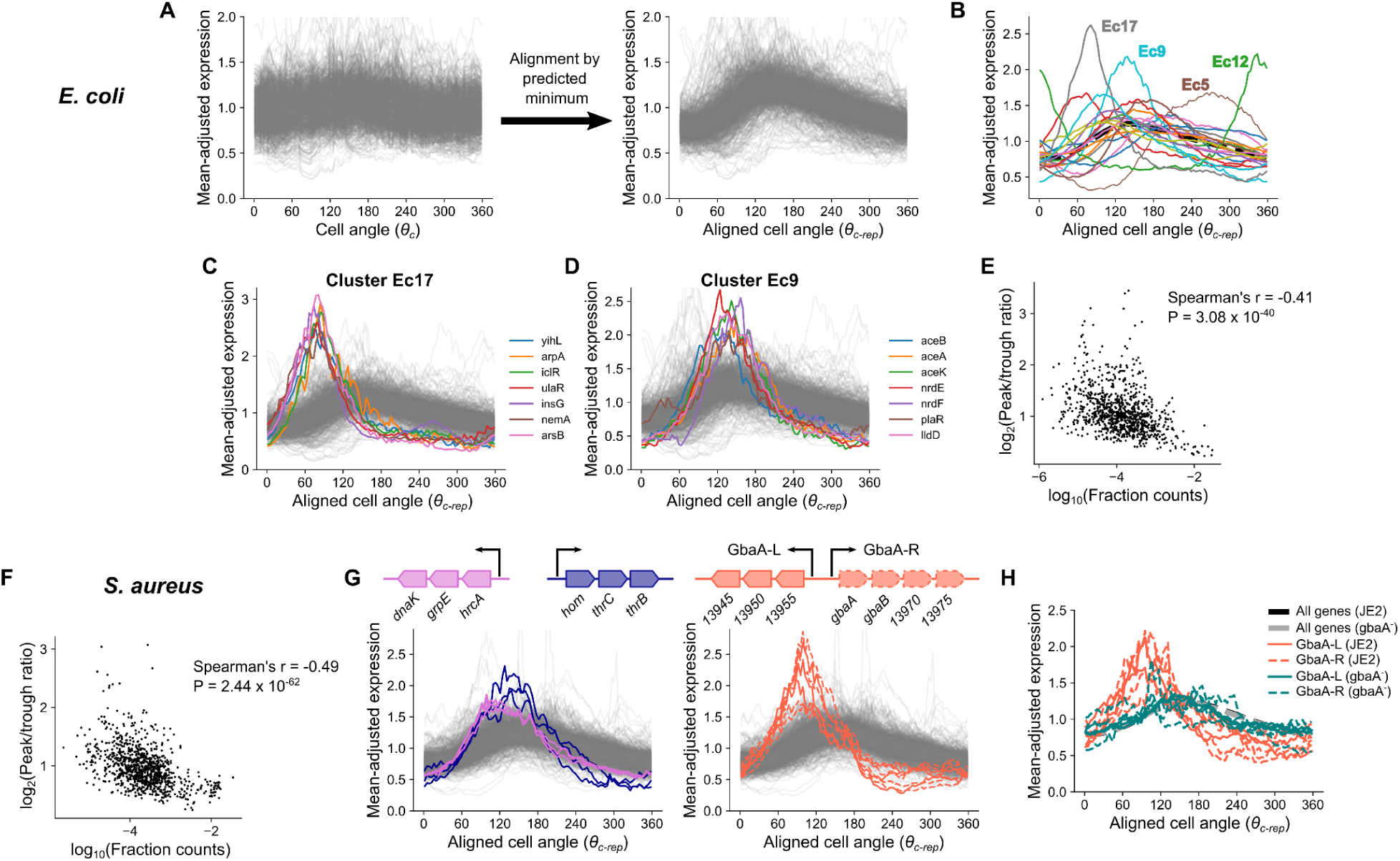
Repression is associated with higher amplitude in cell cycle gene expression. **A)** Procedure to align expression profiles of different genes. Smoothed expression for each gene normalized by division by its mean (*left*) is standardized by rotating cell angle so the predicted replication time expression is at zero. We term this aligned cell angle progression metric *θ_c-rep_*. See Materials & Methods. **B)** Average aligned expression profiles for 20 *k*-means clusters in *E. coli*. The dotted black line represents average expression across all reproducible genes. **C & D)** Plots of individual genes from clusters in **(B)**. **E & F)** Comparison of average expression to the log-ratio of peak to trough expression in *E. coli* **(E)** and *S. aureus* **(F)**. **G)** Aligned expression profiles for select operons in clusters Sa11 and Sa18, with operon structure shown. **H)** Aligned expression profiles for GbaA regulon genes in JE2 and a *gbaA^-^* transposon mutant. Thick black and gray lines represent average expression across all reproducible genes.

To identify the range of behaviors, we partitioned *E. coli* genes into 20 clusters based on the aligned dynamics (Fig. 5B). Of these, several exhibited particularly divergent expression, differing from the expected pattern in both the timing of expression dynamics and the amplitude (i.e. the relative difference between maximal and minimal cell cycle expression). Cluster *E. coli* (Ec) 12 comprised the *nrdAB-yfaE* operon and cluster Ec5 contained the *dnaAN-recF* operon and other delayed expression genes, including some *nuo* genes. Cluster Ec17 showed an early-peaking pulse in expression with greater amplitude than most genes (Fig. 5C). Many genes in these clusters were in operons that encode repressors, at least some of which have autorepressive activity (including *nemA*, which is co-transcribed with the autorepressor *nemR*) (Table S4). Cluster Ec9, whose members peak at the expected time but show increased amplitude (Fig. 5D), also included several repressed genes (Table S4), such as the glyoxylate shunt operon, *aceBAK*, which is IclR-repressed. While these clusters showed the most dramatic patterns, other clusters composed of low-expressed genes showed similar trends (Fig. S14A). Globally, we observed that lower average expression was associated with expression amplitude when amplitude was measured either as peak-to-trough fold change or standard deviation after mean-adjustment (Fig. 5E, Fig. S14C), and this trend was stronger when we focused on only the most-reproducible genes (Fig. S14D & E). Previously, Wang and colleagues^6^ observed that for the *lacZ* gene in *E. coli*, gene replication is associated with a pulse in transcription, but that this effect is reduced as its repression by LacI is relieved. Our data suggest that similar repression-driven effects, while varying greatly between genes, may be present across the *E. coli* transcriptome.

Extending this analysis to *S. aureus*, we also observed a negative relationship between average expression and amplitude of cell cycle expression, suggesting similar principles (Fig. 5F, Fig. S14F & G). After clustering genes based on their aligned dynamics, we noted extreme divergence in several clusters, in which we identified genes belonging to genome-integrated mobile genetic elements (MGEs) (Fig. S15). Genes within these clusters were localized within the core of the MGE, suggesting a role in MGE mobilization as opposed to host-related functions (such as virulence factors)^40–42^. After excluding all MGE genes, however, a range of behaviors were still evident (Fig. S14H). For example, as in *E. coli*, we observed high amplitude and delayed dynamics in a cluster, *S. aureus* (Sa) 9, comprised of *dnaAN*. Analogous to clusters Ec17 and Ec9 in *E. coli*, we observed high-amplitude clusters with (Sa18) and without (Sa11) a “left” shift, indicating that expression peaked earlier than expected (Fig. S14I & J). Sa11 contained a range of genes including the heat shock response operon, *hrcA-grpE-dnaK,* and an amino acid biosynthesis operon, *hom-thrCB*, which showed a particularly large expression amplitude (Fig. 5G). Sa18 was almost exclusively composed of genes in the GbaA regulon (Fig. 5G). In contrast, another cluster (Sa12) showed delayed dynamics (Fig. S14K). Notably, this included several genes involved in stress and virulence.

Since high amplitude in gene expression is typically associated with low average expression levels, and based on previous observations^6,43,44^, we reasoned that transcriptional repression could be driving the high amplitude pulses observed for genes in certain clusters (Ec9, Ec17, Sa11, Sa18). Therefore, we focused on genes of the *S. aureus* GbaA regulon (Fig. 5G), which showed a particularly strong early pulse in expression. This regulon consists of two divergent operons (referred to here as “GbaA-L” and “GbaA-R”) that are repressed by GbaA. GbaA is a transcriptional repressor encoded by *gbaA* within the GbaA-R operon whose repression is relieved by reactive electrophilic species such as quinones or aldehydes^45,46^. To test whether GbaA repression was responsible for the divergent dynamics of its regulon, we compared wild-type expression dynamics to those of a *gbaA* transposon mutant, where GbaA-mediated repression should be relieved. Since transposon insertion happens within the GbaA-R operon, transcription of this locus was disrupted, whereas in the GbaA-L operon we observed a >100-fold increase in expression (Fig. S16A) due to loss of repression. As predicted, this loss of repression was accompanied by a clear reversion of GbaA-L expression to the expected pattern in the transposon mutant, as well as reduced expression amplitude (Fig. 5H). To further verify that this change resulted directly from loss of the regulator rather than disruption of the locus, we measured transcription from the GbaA-L promoter upon integration at an alternative chromosomal locus. While repression by GbaA was less efficient at this locus than for native GbaA-L (Fig. S16B), we nonetheless observed a spike in reporter expression on a wild-type JE2 background that was absent when the reporter was integrated on a *gbaA*^-^ transposon mutant background (Fig. S16C), further supporting that the GbaA regulon dynamics arise due to repressor-promoter interactions. These observations suggest that repression drives the high-amplitude pulses in expression seen for low-expressed genes.

## Discussion

Our analysis reveals, for the first time, the cell cycle transcriptomes of rapidly proliferating bacteria. Although the expression of most genes fluctuates, crucially, these fluctuations do not appear to be a response to cell cycle-dependent changes in the cellular environment (with a few exceptions: DnaA is not only the major regulator of replication initiation^47^, but also regulates its own transcription in a cell cycle-dependent fashion^48,49^, explaining its highly divergent expression in both species). Instead, gene expression fluctuations during the cell cycle appear to be responses to the local perturbation that each gene experiences upon passage of the replication fork. This appears to be the case even for major cell cycle regulators and explains why despite the known cell cycle-dependent fluctuations of *ftsZ*^50,51^,which encodes the major regulator of cell division in *E. coli*, division timing appears to be relatively insensitive to the expression patterns of this protein^52–54^. A direct link between *ftsZ* replication and transcriptional inhibition was previously postulated but the authors at the time could not provide a satisfactory mechanistic explanation^51^. Here, we explain these augmented fluctuations in *ftsZ* abundance as a consequence of transcription from a distant promoter^36^ (Fig. 4B, Fig. S13B). Our observations therefore support the view that the cytoplasm may be relatively invariant during cell cycle progression of bacteria in a state of balanced growth^55^, at least as it pertains to the activity of specific transcriptional modulators. Thus a gene is likely to experience few environmentally-induced changes to its transcription during the cell cycle besides its own replication. While it is important to consider the potential influence of global factors on gene expression (such as competition for RNA polymerase between genes^56,57^), it is not clear which of these could lead to the dynamics we describe here. By redefining cell cycle expression of a gene relative to its replication time, as measured by *θ_c-rep_* (Fig. 5), we explicitly focus instead on the response of each gene after perturbation by its replication. This provides an expression trace specific to each gene, which we here term the Transcription-Replication Interaction Profile (TRIP).

Analysis of each species reveals a diversity of TRIPs that may reflect gene-specific variation in local regulatory motifs. This variation may arise from each gene’s distance from the promoter, local repression state, and possibly other factors such as chromatin structure, together generating a high degree of complexity that we are only beginning to untangle. Nevertheless, we can distinguish several archetypal behaviors of TRIPs (Fig. 6). First, we delineate the non-divergent or “canonical” pattern (Class 1). For genes that fall into this category, expression increases in response to gene dosage at a rate that is likely to be proportional to mRNA half-life^9^, before being gradually diluted as a fraction of total mRNA as gene dosage increases the expression of subsequently-replicated genes. For genes outside this category, we observe divergence of TRIPs along two main axes: *heterochrony*, or differential expression timing, and *heterometry*, or differential amplitude (or “peak/trough ratio”). Many operons under repression exhibit *heterometry* (Class 2 & 3), while a subset of these peak earlier than expected (*heterochrony*) (Class 2). Genes can also exhibit *heterochrony* as a “delayed” expression profile (Class 4). Finally, we note that in *S. aureus*, many genes located in MGEs, particularly those involved in mobilization, exhibit heterogeneity patterns that are entirely distinct from those of the host genome (Class 5). Future work will be required to fully describe the heterogeneous expression of these elements.

**Figure 6:**
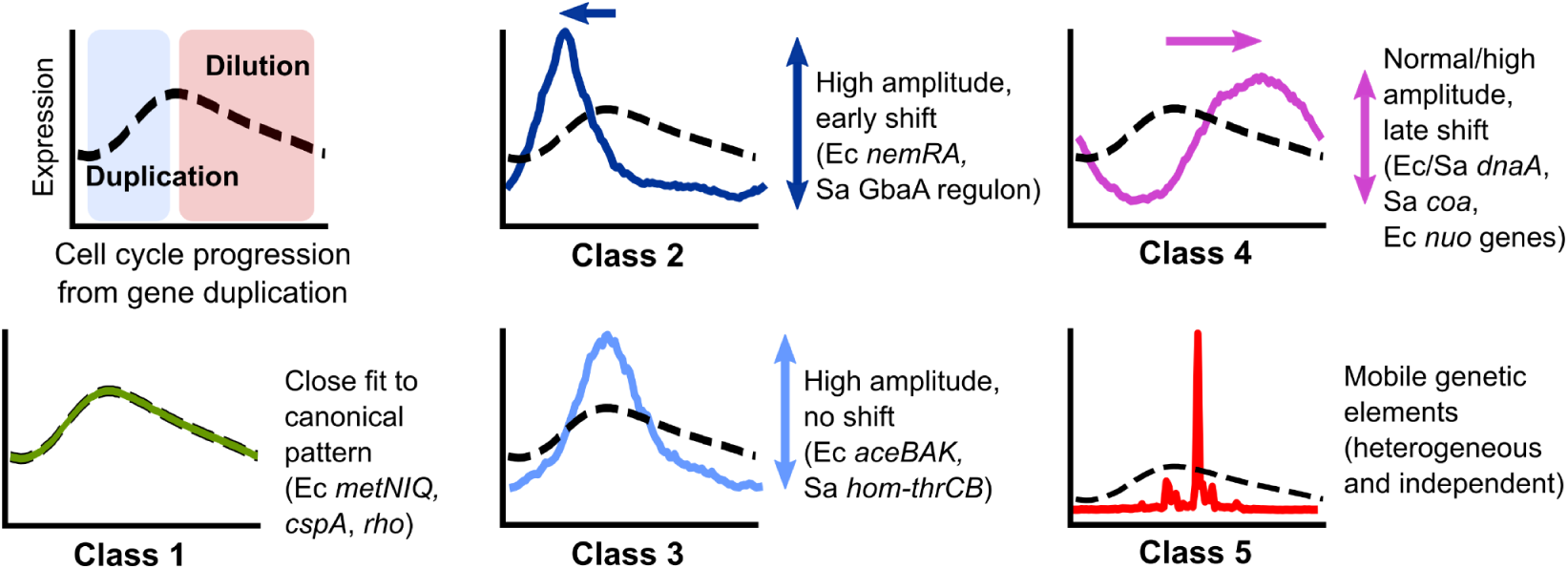
Classes of Transcription-Replication Interaction Profiles of non-divergent and divergent genes. *Top left*: Canonical TRIP driven by gene dosage. *Other panels*: Archetypal patterns of TRIPs that do not (Class 1) or do (Classes 2-5) diverge from this pattern. Genes in *E. coli* and *S. aureus* are represented as Ec and Sa, respectively.

Mechanistically, much remains to be explored. For genes with Class 2 or 3 TRIPs, many genes are under repression (or even autorepression). This suggests a possible mechanism in which the passage of the replication fork through the promoter transiently displaces the repressor, leading to a temporary increase in transcription shortly after replication^6,58^. Other modes of replication-induced transcription have also been suggested^43,44^. However, it is unclear what drives the precise timing of these transient increases. In *E. coli*, *iclR*, which encodes a transcriptional repressor that represses itself as well as the neighboring *aceBAK* operon, has a Class 2 TRIP, whereas its target, *aceBAK*, belongs to Class 3. This demonstrates that the presence of binding sites for a particular repressor may not alone be sufficient to determine the expression timing. For Class 4, the delayed pattern, the effect of gene position within operons in *E. coli* clearly points to the greater disruption experienced by genes far from their promoters, but in other cases, particularly in *S. aureus*, there must be other drivers. Overall, while certain themes emerge, many questions remain about how these myriad influences on gene expression interact to produce the observed patterns.

As our interpretation of these signatures continues to improve, we may be able to distinguish additional modes of regulation. For example, does low expression of a specific gene reflect weak intrinsic promoter strength (subject to positive regulation) or strong repression (subject to negative regulation)? A Class 2 or 3 TRIP would indicate the latter. Alternatively, what does the delay in expression of genes associated with stress responses or virulence in *S. aureus* tell us about their regulation, and how might this relate to the phenotypic heterogeneity in stress sensitivity and virulence observed in bacterial pathogens^59^? Our work demonstrates that this approach can be extended beyond standard model organisms to allow comparison across genes, genetic backgrounds, or even distantly-related species, helping to characterize control of virulence or resistance genes in an emergent pathogen, or regulation of a gene cassette with potential biotechnology applications^60^. Finally, our ability to infer global parameters directly from the data, including replication patterns and both RNA and DNA polymerase speeds, facilitates comparison across very different growth conditions and will allow us to connect gene-specific dynamics to the overall state of the cell.

This work represents only an initial effort in this direction, but provides a foundational framework for genome-wide exploration of novel bacterial regulatory phenomena. As bacterial scRNA-seq methods evolve in scale, capture efficiency, and cost^61–64^, we predict that these methods, in combination with microscopy and molecular genetics approaches that allow mechanistic dissection of these phenomena, will illuminate a diverse ecosystem of dynamic transcriptional processes.

## Materials and Methods

### Bacterial strains and growth conditions

Strains used are listed in Table S1. All *E. coli* strains (a gift from Dr. Christian Rudolph) were routinely grown in modified Luria Broth (LB) (1% tryptone (Sigma-Aldrich), 0.5% yeast extract (Sigma-Aldrich), 0.05% NaCl, pH adjusted to 7.4^22^). For growth in minimal media, an M9 base (1X M9 minimal salts (Gibco), 2 mM MgSO_4_, 0.2 mM CaCl_2_) was supplemented with 0.4% glucose (M9G) or with both 0.4% glucose and 0.2% acid casein peptone (Acros Organics) (M9GA). All *S. aureus* strains were routinely grown in Bacto tryptic soy broth (TSB) (BD Biosciences). The *gbaA* transposon mutant was provided by the Network on Antimicrobial Resistance in *Staphylococcus aureus* (cat. # NR-46898).

### Growth curves

Strains were grown overnight in LB (*E. coli*) or TSB (*S. aureus*) at 37°C, shaking at 225 rpm. For initial experiments with *S. aureus* (Datasets D3 & D4), strains were diluted to an A_600_ value of 0.05 in prewarmed TSB, after which A_600_ was measured at the times specified. A_600_ was measured on a BioMate 3S spectrophotometer (Thermo Scientific). For experiments with *S. aureus* in balanced growth (Datasets D5-D8), overnight cultures were diluted in TSB first to 0.005, then after 3 hr diluted again to 0.005 before measuring A_600_ at the time intervals specified. For *E. coli* growth curves, strains were diluted to an A_600_ value of 0.05 and incubated for 2 hr in the desired medium then diluted again in the same prewarmed medium to an A_600_ value of 0.005, after which A_600_ was measured at the time intervals specified. Where *E. coli* cells were diluted into a different medium, cells were washed once with PBS prior to dilution. To measure growth rate, a linear model log_2_(A_600_) ∼ *m*T + *c* was calculated for the linear portion of this relationship (where T is the time in minutes) using the LINEST function in Microsoft Excel and the doubling time in minutes *t_d_* was calculated as 1/*m*.

### PETRI-seq analysis

Cells were grown as described for the growth curves except that after specific time intervals (for *S. aureus*, 2 hr 20 min in initial experiments, 1 hr 30 min in balanced growth experiments; for *E. coli*, 2 hr, 3 hr, and 7 hr in LB, M9GA, and M9G, respectively, when growth rates appeared constant (Fig. S3)) cells were harvested by centrifugation and resuspension in 4% formaldehyde in PBS. For *S. aureus* initial experiments, centrifugation was at 10,000 x g, 1 min at room temperature and for *E. coli* and balanced growth *S. aureus* experiments, centrifugation was at 3,220 x g, 5 min, 4°C. PETRI-seq was carried out as described previously^16^ with the following modifications. Initial fixing, permeabilization, and DNase treatment were carried out as described but with cell wall permeabilization using 100 µg/ml lysostaphin (Sigma-Aldrich) for *S. aureus* and 100 µg/ml lysozyme (Thermo Scientific) for *E. coli*. For Dataset D4, samples were split into processing with or without DNase treatment and subsequent wash steps, to test whether this would affect correlation patterns (suggesting contaminating genomic DNA could play a role). However, no difference was observed in the presence or absence of DNase treatment, although UMI/barcode was slightly higher after DNase treatment (Table S1). For barcoding, the number of cells included was reduced from 3 x 10^7^ to a maximum of 1 x 10^7^, since preliminary experiments indicated lower input at this stage was associated with a higher UMI/barcode for *S. aureus*. Tagmentation was performed using the EZ-Tn5 transposase (Lucigen) as described in the latest version of the PETRI-seq protocol (available at https://tavazoielab.c2b2.columbia.edu/PETRI-seq/updates_April2021/PETRI_Seq_Protocol.pdf). Briefly, the transposase was loaded by incubating EZ-Tn5 with pre-annealed oligonucleotides (/5Phos/CTGTCTCTTATACACATCT and GTCTCGTGGGCTCGGAGATGTGTATAAGAGACAG) at 4 µM and 40% glycerol at room temperature for 30 min. Tagmentation was then performed incubating samples with loaded EZ-Tn5 (at a final further dilution of 400x) and 2x Tagment DNA buffer; either using the Nextera 2x Tagment DNA (TD) buffer or 20 mM Tris(hydroxymethyl)aminomethane; 10 mM MgCl_2_; 20% (vol/vol) dimethylformamide, pH adjusted to 7.6 with acetic acid^65^. After incubating for 5 min at 55°C and decreasing the temperature to 10°C, either Nextera NT buffer (Illumina) or 0.2% sodium dodecyl sulfate was added, allowing neutralization to proceed for 5 min at room temperature. Final amplification was performed with Q5 polymerase (New England Biolabs) using the NEBNext Universal i5 primer (New England Biolabs) and the N7 indices from the Nextera XT Index Kit v2 Set A (Illumina) as also described in the updated PETRI-seq protocol. Sequencing was performed on an Illumina NextSeq 500 to obtain 58 x 26 base paired-end reads. For each barcoding experiment, multiple libraries of ∼20,000 cells were prepared and sequenced, and no batch effects were noted across libraries.

### Pre-processing and scVI analysis

Initial demultiplexing of barcodes, alignment, and feature quantification was performed using the analysis pipeline described in ^16^ except that feature quantification was performed at the gene level rather than operon level. Reference sequences and annotations were obtained from Genbank (https://www.ncbi.nlm.nih.gov/genbank/). *E. coli* reads were aligned to the K-12 MG1655 reference assembly (GCA_000005845.2) and *S. aureus* to the USA300_FPR3757 reference assembly (GCF_000013465.1). After initial processing, counts by cell barcode were pooled across different libraries and initial filtering was performed using Scanpy v1.7.1^66^. Barcodes with UMI below a threshold (15 for Dataset D1, D2, D4; 20 for Dataset D3, D5-7, 40 for Dataset D8) were removed, as well as any genes with fewer than 50 UMI across all included barcodes (100 for Dataset D3). To generate the denoised representation of the data, scVI v0.9.0^20^ was applied with the following hyperparameters, chosen through grid search to distinguish between closely related *S. aureus* strains in a pilot dataset: two hidden layers, 64 nodes per layer, five latent variables, a dropout rate of 0.1, and with a zero-inflated negative binomial gene likelihood (other hyperparameters maintained as defaults). Denoised expression values based on the scVI model were obtained using the scVI function “get_normalized_expression”.

### Cell cycle analysis

Cells were assigned to cell cycle phases by calculating the angle *θ_c_* relative to the origin between *x* and *y* coordinates in a two-dimensional UMAP embedding of the data as tan^-1^(x / y), similar to the ZAVIT method our lab has described previously^67,68^. scVI-denoised expression values were first log_2_-transformed then converted to *z*-scores. Embeddings were computed by averaging these *z*-scores within bins according to chromosomal location (50-400 kb bins, depending on the dataset), and then performing two-dimensional UMAP analysis using the umap-learn v0.5.1 library in Python (https://umap-learn.readthedocs.io/en/latest/) with the ‘correlation’ distance metric. These embeddings were then mean-centered (Fig. 2A & Fig. S7B). To get the expression by cell angle matrix used in Fig. 2B, gene expression *z*-scores were then averaged within 100 equally spaced bins of *θ_c_* to produce a cell angle-binned expression matrix. To order genes based on their cell cycle expression, gene angle, *θ_g_*, was calculated as follows. PCA was performed on the transpose of the cell angle-binned expression matrix and *θ_g_* was calculated as the angle between PCs 1 and 2 relative to the origin. Together, *θ_c_* and *θ_g_* are metrics for ordering of cells and genes, respectively, within the model of cell cycle gene expression described here.

### Modeling the gene angle-origin distance relationship

While there was a strong relationship between origin distance *D* and gene angle *θ_g_*, modeling this relationship is challenged by the fact that the relationship is “wrapped” with an unknown periodicity with respect to *D* (Fig. 2E & F, Fig. S7D) (i.e. after a period of increased *θ_g_* with *D*, *θ_g_* starts again at zero). To fit this relationship, a custom Bayesian regression analysis was developed according to the following model partially adapted from ^69^, with both *θ_g_* and *D* standardized to the range -π to π:

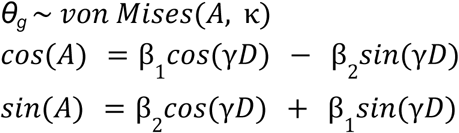

Where:

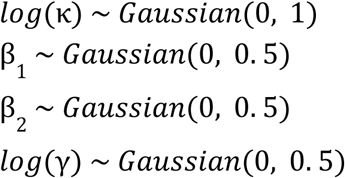

The von Mises probability distribution is a circular probability distribution here parameterized by *A*, the predicted mean angle, and *κ*, the concentration parameter (higher κ implies greater concentration of the distribution around *A*). The parameter *ɣ* can be interpreted as the gradient of *D* with respect to *θ_g_* after standardizing both variables to to the range -π to π. The inverse of *ɣ*, 1/*ɣ*, is the gradient of *θ_g_* with respect to *D* (after range standardization) and therefore is the fraction of the origin-terminus distance covered within a single span of *θ_g_*. Therefore, 1 - 1/*ɣ* is the fraction of *D* during which the next round of replication has already initiated, referred to as the “overlap fraction” in Fig. 2G & Fig. S7E. Here, *ɣ* is constrained to be positive by the lognormal prior distribution (Fig. S17), which is appropriate since the ordering of angles *θ_g_*are reversed (i.e. 360 - *θ_g_* when *θ_g_* is in degrees) if during analysis this relationship shows a negative trend. This can occur because the directionality of PCs used to calculate *θ_g_* is arbitrary. Posterior distributions for the parameters were obtained by Hamiltonian Monte-Carlo sampling using Rstan v2.21.3^70^. Fitted values for *θ_g_*based on *D* (*θ_g-pred_*) were calculated by determining *θ_g-pred_* for all sampled parameter values and then calculating the mean value of *θ_g-pred_* as tan^-1^(mean(sin(*θ_g-pred_*)) / mean(cos(*θ_g-pred_*))).

#### Calculating replication pattern statistics

We can use the gradient parameter, *ɣ*, of the gene angle-origin distance model to calculate statistics of the replication pattern. The parameter *ɣ* can be interpreted as the gradient of *D* with respect to *θ_g_* after standardizing both variables to to the range -π to π. To convert the gradient to °/Mb (as in Fig. S7F), this value is multiplied by 360 divided by origin-terminus distance in Mb. The average DNA polymerase speed can be estimated from this as follows:

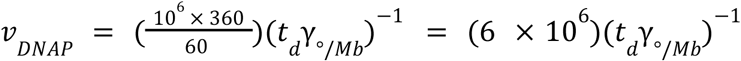

Here, *v_DNAP_* is the DNAP speed in bp/s, *t_d_*is the doubling time in min, *ɣ_°/Mb_* is the gradient of the gene angle-origin distance relationship in °/Mb.

### Modeling the cell angle-gene angle relationship

To predict expression based on cell angle *θ_c_* and gene angle *θ_g_*, a linear regression model was constructed using scikit-learn v0.24.1^71^ with features generated from *θ_c_* and *θ_g_*. Specifically, both angles were converted to radians and then transformed into cos(*θ_c_*), sin(*θ_c_*), cos(*θ_g_*), and sin(*θ_g_*). All interactions and combinations of these terms up to a fourth degree polynomial were constructed using the scikit-learn PolynomialFeatures function. The untransformed *θ_c_* and *θ_g_*values in radians were also included as features. These features were then used to fit a Ridge regression model (ɑ = 10). The model was trained on scVI expression *z* scores averaged first in 100 bins by *θ_c_* then in 100 bins by *θ_g_* (i.e. the expression matrix used for Fig. 3F). An alternative approach considered was a non-linear approach using the scikit-learn implementation of kernel ridge regression with kernel “rbf”. However, the fourth degree polynomial model performed similarly and was computationally far more efficient so was chosen (increasing the polynomial degree further made little difference to performance).

### Predicting expression dynamics based on DNA replication alone

To derive a prediction of cell cycle gene expression dynamics based on the expected effect of replication alone, the two regression models above were combined to yield the pipeline in Fig. S8. Firstly, the gene angle-origin distance model (see Section “Modeling the gene angle-origin distance relationship”) was used to predict the expected value *θ_g-pred_* from origin distance *D.* Next, cell cycle expression was predicted using the cell angle-gene angle regression model (see Section “Modeling the cell angle-gene angle relationship”) using *θ_g-pred_*values. For cell angle *θ_c_*, values used were the average *θ_c_*values of cells binned into 100 equally spaced bins by *θ_c_*. This gives a replication-predicted gene expression matrix of 100 bins x number of genes. The success of this model fit was evaluated based on the correlation with the *θ_c_*-binned expression *z*-scores derived from scVI (Fig. S9A & F), as well as the loss of global chromosome position-dependent gene-gene correlations upon correction of scVI expression with replication-predicted expression (Fig. S9B & G). Additionally, we used this modeling approach to set the zero angle for gene expression plots.

#### Setting the position of *θ_c_* = 0

Initially, the cell angle *θ_c_* orders cells by their cell cycle position within a circle but the start point, when *θ_c_*= 0, is arbitrary. This is not only challenging to interpret but impedes comparing across replicates. Therefore, we standardized *θ_c_* so that *θ_c_* = 0 was the predicted point of replication initiation. Using the inference approach described above, we predicted the gene expression profile by *θ_c_* for an imaginary gene at *D* = 0 (i.e. at the origin of replication). We then determined the value of *θ_c_* giving the minimum predicted expression, reasoning that if increased expression in this model is responsive to a doubling of copy number, the doubling event should occur at the expression minimum. Therefore, we determined this angle, *θ_0_* to be the most likely value of *θ_c_* at which replication initiation occurs, rotating the angles by the operation (*θ_c_* - *θ_0_*) *mod* 360 to set this point as 0°. This interpretation is roughly in accordance with the estimated timing of replication initiation as determined directly from smFISH data (Fig. S10F and see Section “Inferring cell-cycle phase from the DAPI signal”). Crucially, however, it also provides a point of standardization that allows in-phase comparison of cell cycle expression profiles across independent replicates.

### Identifying replication-divergent genes

We identified replication-divergent genes based on two criteria: absolute variability by cell angle *θ_c_* and divergence from the replication model.

#### Identifying genes with high cell cycle variance

First, we identified highly variable genes as follows (based on the method implemented in Seurat v3^72^). We normalized raw counts for library size (so that the total sum of UMI for each barcode was the median UMI/barcode), then to reduce sparsity while retaining cycle information, we averaged counts across 20 bins by *θ_c_*. Next, we log_2_-transformed the data (removing any genes with zero values after binning to allow log-transformation). We observed a negative overall relationship between the mean and variance of genes in log-transformed data (Fig. S9C), to which we fitted a regression line with locally weighted scatterplot smoothing (LOWESS) using the Python package statsmodels v0.12.2^73^. We used this fit to develop a mean-dependent variance threshold. In all cases, genes were considered highly variable if they had a ratio of observed to LOWESS-predicted variance > 1.3 as well as a log_2_ mean normalized expression > −10. These thresholds typically classified ∼25% of genes as highly variable.

#### Identifying genes with high divergence from predicted expression

Next, to quantify divergence from the replication model, we subtracted the replication-predicted expression from the scVI-derived expression *z*-scores (both averaged in 100 bins by *θ_c_*) to “correct” for the effect of replication, and then calculated the standard deviation of this replication-corrected value, *σ_corrected_*. A high *σ_corrected_* indicates that the dynamics behave differently from that expected based on replication alone. Thresholds for *σ_corrected_* (0.6 for *E. coli*, 0.5 for *S. aureus*) were determined manually based on inspection of the relationship between *σ_corrected_* across two datasets and choosing a value above which the correlation between datasets was stronger (Fig. S9E & I) (below the threshold, lack of reproducibility of *σ_corrected_*suggests divergences are small and dominated by noise). To calculate peak/trough fold changes in expression, normalized gene expression derived from scVI was averaged into 100 bins by *θ_c_* and then the ratio between the fourth highest and fourth lowest values were calculated (this was chosen instead of maximum/minimum values to increase robustness to noise).

### Analyzing the effect of operon gene position on expression dynamics

We identified the excess of genes with a “delayed” expression profile by calculating the angle difference as *tan^-^*^1^*(sin(θ_g_ – θ_g-pred_) / cos(θ_g_ – θ_g-pred_))* where *θ_g_* and *θ_g-pred_* are the observed and predicted gene angles in radians, respectively. For operon annotations, *E. coli* transcription units from Biocyc ^74,75^ (https://biocyc.org/) were used. To investigate the relationship between gene distance from transcriptional start sites and angle difference in *E. coli*, all genes in polycistrons (transcription units with more than one gene) were included. The distance was measured from the annotated transcription unit start site to the midpoint of each gene. Where genes were in multiple transcription units, the longest distance from a start site was taken. Angle difference was converted into time by dividing the angle by 360° then multiplying by the doubling time in seconds. For *S. aureus*, operon annotation was obtained from AureoWiki^76^ (aureowiki.med.uni-greifswald.de). Since this provided only the genes within an operon and not its start, the first base of the first gene was taken as the transcriptional start site.

#### Per-base analysis of the *nuo* and *mraZ-ftsZ* operons

To analyze per-nucleotide coverage of the *nuo* operon (Fig. 4D & E), we obtained “.bam” alignment files from the analysis pipeline (see “Pre-processing and scVI analysis) and removed PCR duplicates with UMI-tools v0.5.5^77^. Next, for a genomic interval encompassing the *nuo* operon and neighboring genes, we quantified per-base per-barcode read depth using the *mpileup* function in Samtools v1.3.1^78^. This coverage was then normalized by total per-cell library depth (division by per-cell total mRNA count then multiplication by median mRNA count across all cells) and averaged in 10 bins by *θ_c_*. For the plots in Fig. 4D & E, we recenter *θ_c_* so that 0° is the predicted minimum expression of *nuoA*, the first gene in the operon, so that *θ_c_* corresponds to the approximate time elapsed since the locus was replicated. Analysis of the *mraZ-ftsZ* locus was carried out as for the *nuo* operon except that *θ_c_*was recentered so that 0° is the predicted minimum expression of *mraZ*.

### Aligning gene expression profiles of based on their predicted minimum expression

To align cell cycle gene expression profiles as displayed in Fig. 5A & C, we use the replication-predicted expression profiles derived above to determine the minimum cell angle, *θ_c-min_*, predicted for each gene. Profiles of gene expression by cell angle (averaged in 100 bins by *θ_c_* as used elsewhere) are then rotated so that *θ_c_*= 0 corresponds to this new minimum by the transformation (*θ_c_* - *θ_c-min_) mod* 360 to give the cell angle relative to the predicted timing of a gene (*θ_c-rep_*). Gene expression profiles are then divided by their mean to center them, but they are not scaled (so that amplitude differences are preserved). These profiles are used to generate the *k*-means clusters described.

### Simulating the effect of DNA replication on gene expression

We predicted the gene-gene correlation patterns arising from DNA replication using a simulation written in Python (see Fig. S4) as follows. Cells were represented by genomes with 200 genes, each represented as a single integer and divided into individual replication units. In the simplest case, genomes were divided into two units of 100 genes (i.e. the two “arms” of the chromosome). In each cell, replication initiation events were simulated at intervals determined by a Poisson distribution with expected value μ. After an initiation event, replication proceeds in stepwise fashion along the length of each replication unit, doubling the copy number at each point until the end of that replication unit has been reached. We also simulate “cell division” events in which all copy numbers are halved. These are timed independently from replication initiation but in the same way (at Poisson-distributed intervals with rate μ), with an additional offset from the first replication initiation event. In practice, we found that this offset did not affect correlations, since all genes are scaled equally. We used an initial offset of 150 steps (i.e. 1.5x the time to replicate a 100 gene replication unit, equivalent to the 40 min C-period + 20 min D-period originally proposed for *E. coli* B/r ^4^). For each simulation, we generated 1,000 cells. Cells were initiated one at a time to yield an unsynchronized population, then the simulation was run for a further 1,000 steps with the whole population. We then normalized expression by total counts and calculated Spearman correlations across all genes. In order to simulate specific doubling times, the rate μ was calculated as *μ = (n* × *t_d_) / t_c_* where *n* is the number of genes in the longest replication unit (here, 100 genes), *t_d_* is the doubling time, and *t_c_* is the C-period (here a value of 42 min was chosen for *E. coli* MG1655 based on ^79^). The *t_d_*/*t_c_*ratio represents the fraction of one round of chromosomal replication that can take place in one cell cycle. Finally, for simulation of cells with additional origins of replication, genes were split into replication units according to the following assumptions: a) all origins initiate replication simultaneously; b) replication stops at the termination site *ter*, which is halfway along the chromosome; c) genes are replicated by the nearest origin (unless the replication fork must pass through *ter* to reach that gene).

### Bulk RNA-seq analysis

For the analysis of bulk RNA-seq from ^11^ (Fig. S4C), we accessed data from the Gene Expression Omnibus (GEO, https://www.ncbi.nlm.nih.gov/geo/) under accession ID GSE46915. Counts were size factor-normalized with DESeq2 v1.32.0 ^80^, then data were standardized to *z-*scores and averaged into 100 kb bins by chromosomal position. Spearman correlations of binned values across all time points and replicates are shown.

### Single-molecule fluorescence in situ hybridization (smFISH)

Our smFISH protocol was described previously^30,81^. Briefly, we first designed seven sets of antisense DNA oligonucleotide probes. Six probe sets were against *E. coli* mRNAs *dnaA*, *nrdA*, *nemA*, *metN*, *rho*, and *cspA*, and another against bacteriophage lambda *cI* mRNA (which serves as a negative control, since the probes have no target in the bacterial cell). All oligos were synthesized with a 3’ amine modification (LGC Biosearch Technologies). The oligos against a given gene (oligo set) were pooled and covalently linked to 5-Carboxytetramethylrhodamine succinimidyl ester (5’-TAMRA SE, Cayman Chemical) and purified using ethanol precipitation. Probe sequences are listed in Table S2.

### Microscopy

An inverted microscope (Eclipse Ti2E, Nikon), equipped with motorized stage control (TI2-S-SE-E, Nikon), a universal specimen holder, an LED lamp (X-Cite XYLIS), a CMOS camera (Prime 95B, Photometrics), and a ×100, NA 1.45, oil-immersion phase-contrast objective (CFI60 Plan Apo, Nikon) was used for imaging. The following fluorescent filter sets were used: DAPI (Nikon, 96370) and Cy3 (Nikon, 96374).

*E. coli* cells were grown as described in Section “Bacterial strains and growth conditions”. After overnight culture, dilution, and re-dilution at 37°C, 220 rpm, cells were grown to a density of ≈ 0.2, then for each gene, 36 ml of culture was collected, immediately fixed and permeabilized, then incubated with the fluorescent probe set, washed. Next, we loaded 2 μl of the cell suspension on a circular coverslip, then covered it by a 1 × 1 cm agarose pad made of 1.5% agarose (Sigma) in 1× PBS, as described in ^30^. The coverslip was then lodged in an Attofluor Cell Chamber (Invitrogen), which was then placed onto the microscope’s slide holder and the cells were visually located using the phase-contrast channel. Images were taken in the following order: phase-contrast (100 ms; to detect the cell outline), Cy3 (400 ms; smFISH-labeled mRNA), and DAPI (4′,6-diamidino-2-phenylindole) (100 ms; bacterial DNA). Snapshots were taken at seven z-positions (focal planes) with steps of 300 nm. Images were acquired at multiple positions on the slide, to image a total of 500–2000 cells per sample (typically 9-16 positions).

### Cell segmentation

Cells were identified in the phase-contrast channel, as described previously^6,81^. Briefly, we first defined the “in-focus” z-slice in every image stack by finding the one with the highest variance among pixels. We then used U-Net, a convolutional network for image segmentation^82^, previously trained on our *E. coli* images, to recognize all pixels that are within any given cell. Finally, the segmentation results were manually inspected, with poorly segmented cells manually corrected or removed.

To estimate the dimensions of each cell, the cell area *A* was first measured by counting the number of pixels within the cell, and the cell length *L* by calculating the length of its long axis. Approximating the bacterial cell as a spherocylinder^83^, we estimated the cell width *d* and cell volume *V* using the equations below:

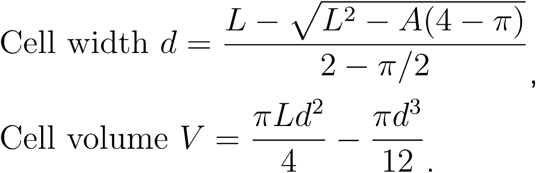

The estimated cell volume *V* is used when measuring mRNA concentrations in each cell (Section “mRNA quantification”), and the cell length *L* serves as an indicator for cell cycle progression (Section “Cell-cycle analysis of smFISH data”).

### mRNA quantification

Following cell segmentation (Section “Cell segmentation”), we estimated the mRNA copy number in individual cells using two methods: (i) based on the recognition of fluorescent foci (“spots”), and (ii) based on the measurement of whole-cell fluorescence. The two methods yielded consistent results (Fig. S12) and were used interchangeably in subsequent analysis.

#### Spot based quantification

Spot recognition and the subsequent mRNA quantification were done as described previously^30,81^. Briefly, we used the Spätzcells software^30^ to identify the spots in the fluorescent images. The software fits the fluorescence intensity profile near each spot to a two-dimensional elliptical Gaussian. The fitting results yielded the properties of each spot, including the position, spot area, peak height (amplitude of the fitted Gaussian), and spot intensity (integrated volume under the fitted Gaussian), used in the subsequent analysis.

To discard false positive spots, such as the ones resulting from nonspecific binding of smFISH probes, we performed a gating procedure as described in ^30,81^. Briefly, we compared the 2D scatter plots of peak height versus spot area for all detected spots in the experimental samples to that from the negative control (the sample incubated with probes against lambda *cI*, see Section “smFISH”). We then defined a polygon in the 2D plane, such that most spots from the negative sample were located outside of it. All spots outside of this polygon were discarded, and the gating results were confirmed by manual inspection of a subset of images.

Following spot recognition, we estimated the fluorescence intensity of a single mRNA molecule as described in ^30^. We fitted the histogram of spot intensities in each experimental sample to a sum of three Gaussians corresponding to one, two, and three mRNA molecules per spot. The center of the first Gaussian was then used to estimate the fluorescence intensity of a single mRNA molecule. Using this procedure, we found that the Gaussian fitting results for genes *dnaA*, *nrdA*, *nemA*, *metN*, and *rho* were very close to each other, consistent to the fact that the probe sets against them have the same number of probes (see Table S2). Therefore, we used the mean of their first-Gaussian center as our estimated single-mRNA intensity. The high expression level of the *cspA* samples (Fig. S10B) was likely to hinder the identification of individual mRNA molecules^30^. Since the number of probes in the *cspA* set is 1/3 of that against other genes (Table S2), we assumed its single-mRNA intensity to be a third of that for the other genes. Finally, the mRNA copy number for a given gene in each cell was calculated by summing the mRNA spot intensities within the cell and dividing by the single-mRNA intensity^30^, and the mRNA concentration for a given gene in each cell was calculated by dividing the mRNA copy number by the estimated cell volume (Section “cell segmentation”).

#### Whole-cell based mRNA quantification

An alternative approach to relying on spot recognition is the use of total cell fluorescence as a proxy for the total number of bound probes, in turn indicating the number of target mRNA molecules. We first chose the z-slice with the largest coefficient of variation among intracellular pixels, indicating maximum contrast. Next, we determined the background fluorescence intensity by calculating the average fluorescence per intracellular pixel in the negative control (the sample incubated with probes against lambda *cI*, see Section “smFISH”). After subtracting this background intensity from cells in each positive sample, we calculated the total and average (per pixel) fluorescence of each cell. These values exhibited a linear relation with the spot-based measurements of mRNA number and concentration, respectively (Fig. S12). The fitted slopes were used as calibration factors to convert the whole-cell fluorescent signals to mRNA numbers and concentrations.

### Modeling the distribution of cell length

Within a population of exponentially growing cells, under the assumption that the instantaneous growth rate a cell is proportional its length, the cell length distribution is predicted to follow^84^:

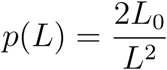

with *L*_0_ the cell length at birth. To account for the stochasticity of cell-cycle processes^85^, as well as the experimental error, we described the measured cell length data using a Gaussian-smoothed version of the original function:

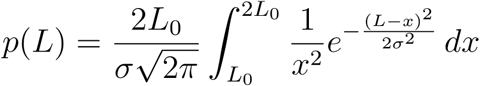

where σ represents the noise magnitude. Fitting this equation to the experimental data (Fig. S10C) yielded *L*_0_ = 3.43 ± 0.05 μm, σ = 0.56 ± 0.10 μm (N = 12 samples, each with > 500 cells. See Table S3 for detailed sample sizes).

### Cell-cycle analysis of mRNA concentration

Comparing the mean expression levels of the six genes (*dnaA*, *nrdA*, *nemA*, *metN*, *rho*, and *cspA*) as measured by smFISH with the estimated abundance obtained by scRNA-seq showed that the two methods were highly correlated(Fig. S10D). We next aimed to test whether the cell-cycle dependence of transcription, revealed by scRNA-seq (Fig. 3 B & D, 2nd column) is too found in the smFISH data.

We first examined the cell cycle dependence of mRNA concentration, since we reasoned that those values would correspond closely to the mRNA fraction measured in scRNA-seq. For this purpose, we followed the approach of ^6^ and used cell length as an indicator for cell cycle progression. In each sample, we first found the two-fold range of cell length containing most cells. The lower bound of this range provides an estimate for the cell length at birth (*L*_0_), and the value found (*L*_0_ = 3.34 ± 0.07 μm, N = 12) was consistent with the estimate in Section “Modeling the distribution of cell length”. The measured single-cell mRNA concentration was binned based on cell length (with each bin containing 10% of the cells in the sample, and a shift of 1 cell between adjacent bins), and the average mRNA concentration within each bin was calculated (Fig. 3 B & D, 3rd column). For all genes, we observed that the mRNA concentration fluctuates along the cell cycle, returning at cell division (length of 2*L*_0_) to a level similar to that at cell birth (length of *L*_0_), as expected.

To directly compare cell cycle patterns between smFISH and scRNA-seq, we needed to correct for differences in both amplitude and phase of the two signals. In particular, whereas the smFISH pattern is aligned by cell length, hence the bacterial birth-to-division cycle, the scRNA-seq data is aligned, through the cell angle, to the timing of genome replication (*oriC* replication to next *oriC* replication). Aligning the two signals was done as follows. We first linearly converted the cell length to a parameter *β* within the range 0 to 2π:

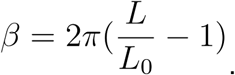

Next, we fitted the relationship between smFISH-measured mRNA concentration and *β* to a sinusoid:

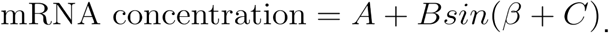

In this function, *A* and *B* indicate the median level and fluctuation of the mRNA concentration, and *C* indicates the phase. Specifically, the maximal mRNA concentration is reached when 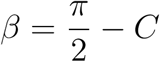 or 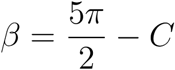 (Fig. S10E).

Similarly, for the scRNA-seq data, we fit the relationship between the mRNA fraction and cell angle *θ_c_* to a sinusoid:

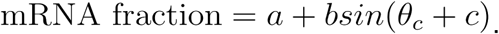

We then estimated the cell angle at cell birth using the phase difference *Φ = C – c* between the fits for scRNA-seq and smFISH data (Fig. S10E). This estimated value (∼155°) was consistent across the 6 genes examined (Fig. S10E).

To overlay the scRNA-seq and the smFISH data (Fig. 3B & D, 4th column and Fig. S11), we scaled and shifted the measured values using the fitting parameters above. The experimentally measured mRNA concentration (smFISH) and fraction (scRNA-seq) were converted using the equations below:

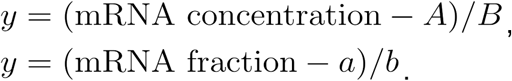

The cell angle *θ_c_* was first shifted by the estimated phase difference, then linearly converted to the corresponding cell length using the equations below:

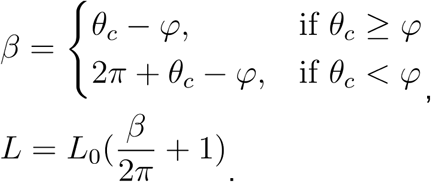

Specifically, the cell length at which *oriC* replicates is estimated to be

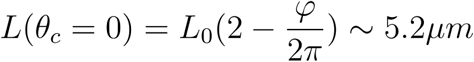

### Comparison to a replication-transcription model

In the simplest model of cell cycle dependent transcription, mRNA levels follow gene dosage, and will thus double following gene replication. To test whether the non-divergent patterns (revealed by scRNA-seq) correspond to this simple scenario, we first binned the smFISH-measured mRNA numbers based on cell length (each bin contains 5% cells in the sample, with a shift of 1 cell between adjacent bins) (Fig. 3B & D, 5th column). Following ^6^, we then fitted the data to the sum of two Hill functions, corresponding to two gene replication rounds:

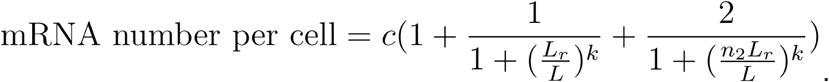

In this expression, the parameter *L_r_* indicates the cell length at which gene replication occurred, and *n*_2_ indicates the fold change in cell length between successive replication events. As seen in Fig. 3B & D, 5th column, the data for the three genes defined as non-divergent (*metN*, *rho*, *cspA*) is well described by this expression, with the fitted *n*_2_ close to 2 as expected (*n*_2_ = 1.89, 2.04, and 2.04 respectively for *metN*, *rho*, and *cspA*). In contrast, two of the three divergent genes (*dnaA* and *nrdA*) exhibit a noticeable deviation from the expected form. In particular, mRNA levels appear to overshoot, consistent with our previous observation^6^.

### Inferring cell-cycle phase from the DAPI signal

When comparing the cell cycle expression patterns obtained by scRNA-seq and smFISH (Section “Cell-cycle analysis of mRNA concentration”), we aligned the two datasets by horizontally shifting by a constant cell-length interval of ∼1.4 μm, equivalent to cell angle of ∼155° (Fig. S10E). This shift is interpreted as corresponding to the cell cycle interval between cell birth and *oriC* replication (which was estimated to take place at cell length of ∼5.2 μm). Whereas in Section “Cell-cycle analysis of mRNA concentration” this value was inferred directly from the mRNA data, we also attempted to estimate the same parameter from single-cell measurements of DNA contents in the smFISH samples, obtained using DAPI labeling (Section “Microscopy”).

We assume that the replication speed is constant along the genome, and designate by *T, T_C_, T_D_* the cell doubling time, duration of genome replication, and the time between replication termination to cell division^79^. We specifically consider the case *max(T_D_, T*/2 *We assume that the repli T < (T_C_ + T_D_*/2) where genome replication initiates at cell age 3*T – T_c_ – T_D_* ^86^. Under these assumptions, the cellular DNA contents (in equivalent number of chromosomes) as a function of cell length (assuming cell length grows exponentially with time^84^, will be given by^86^:

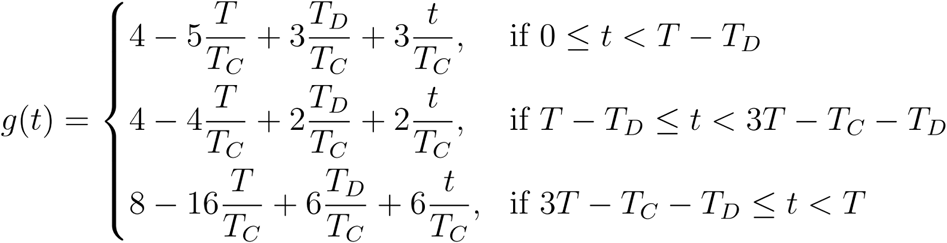

*T – T_D_* is the cell age when one round of genome replication ends, and 3*T – T_C_ – T_D_* is the cell age when another round of genome replication begins. When *t < T – T_D_*, there are three pairs of replication forks present. When *T – T_D_ ≤ t <* 3*T – T_C_ – T_D_*, there are only two pairs of replication forks. When *t* ≥ 3*T – T_C_ – T_D_*, there are six pairs of replication forks. Therefore, the ratios of DNA production rates during these three phases are 3:2:6 (Fig. 10F). In particular, a 3-fold jump in slope takes place at the cell cycle age (length) when *oriC* replicates. We use this constraint to fit our experimental data. We first plotted the single-cell DAPI fluorescence against cell length. We then determined the two-fold range of cell length containing most cells (see Section “cell-cycle analysis of mRNA concentration”), and fitted the data within this length range to the equation above. Discarding those fits where the fitted parameters fell on the boundary of the allowable range and whose r-square value was less than 0.4, the average fitted cell length when the replication of *oriC* occurs is 4.0 ± 0.3 μm (N = 6, with 6 samples discarded). The imperfect agreement between this estimate and the one obtained from scRNA-seq/smFISH alignment (5.2 µm) reflects multiple sources of error. Most notably, the analyses above assumed a simple linear mapping from both cell angle (scRNA-seq) and cell length (smFISH) to cell age, but the relation between observables is in fact nonlinear and subject to stochastic effects. These conceptual errors are likely compounded by experimental ones, for example, the distortion of cell length during fixation, and heterogeneity in DAPI staining.

### Generation of chromosome-integrated reporter constructs in S. aureus

For generation of the reporter construct, we modified the pJC1111 vector^87^, which integrates at the SaPI1 chromosomal attachment (*att_C_*) site. The vector was linearized with restriction enzymes SphI and XbaI (New England Biolabs) and insertion fragments were amplified using Q5 polymerase (New England Biolabs). For the GbaA-L promoter, the intergenic region of the GbaA regulon (130 bp upstream of the *SAUSA300_RS13955* start codon) amplified from USA300 LAC genomic DNA using primers 5’-CCGTATTACCGCCTTTGAGTGAGCTGGCGGCCGCTGCATGGATTACACCTACTTAAAATTCTCTAAAATTGACAAACGG-3’ and 5’-AGTTCTTCTCCTTTGCTCATTATCAACACTCTTTTCTTTTATGATATTTAATAGTTATTGCAAATTCA-3’. *S. aureus* codon-optimized sGFP was amplified from the genomic DNA of *S. aureus* USA300 LAC previously transformed with the pOS1 plasmid (VJT67.63^88^) using primers 5’-AAAAGAAAAGAGTGTTGATAATGAGCAAAGGAGAAGAACTTTTCACTG-3’ and 5’-ATAGGCGCGCCTGAATTCGAGCTCGGTACCCGGGGATCCTTTAGTGGTGGTGGTGGTGGTGGG-3’. Fragments were assembled using the NEBuilder HiFi assembly kit (New England Biolabs) and transformed into competent *E. coli* DH5ɑ (New England Biolabs). The plasmid was purified and then electroporated into RN9011 (RN4220 with pRN7023, a CmR shuttle vector containing SaPI1 integrase), and positive chromosomal integrants were selected with 0.1 mM CdCl_2_. Finally, this strain was lysed using bacteriophage 80ɑ and the lysate was used to transduce JE2 and JE2 *gbaA*^-^ strains, selecting for transduction on 0.3 mM CdCl_2_.

## Acknowledgements

We thank Yitzhak Pilpel, Timothée Lionnet, and Fanny Matheis for critical discussions on the project and the manuscript, and Saeed Tavazoie, Sydney Blattman, and Wenyan Jiang for initial advice on implementing PETRI-seq. We thank Christian Rudolph and his lab for providing the *E. coli* strains. We also further thank Menyu Wang and members of the Yanai and Golding labs for advice and suggestions. The following funding was provided by the National Institutes of Health: R21AI169350 (IY), R01AI143290 (IY), R01AI137336 (BS, VJT, IY), R35 GM140709 (IG).

---

**Figure S1:**
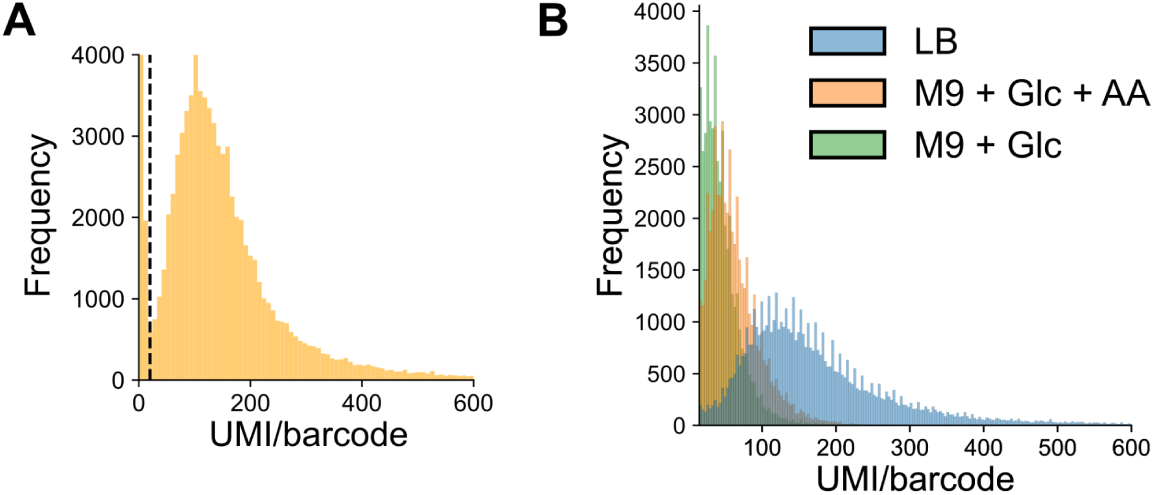
mRNAs captured per cell by PETRI-seq. mRNA captured is quantified as unique molecular identifiers (UMI) per unique cell barcode combination. **A)** *S. aureus* in TSB from Dataset D3. **B)** *E. coli* in different media from Dataset D1.

**Figure S2:**
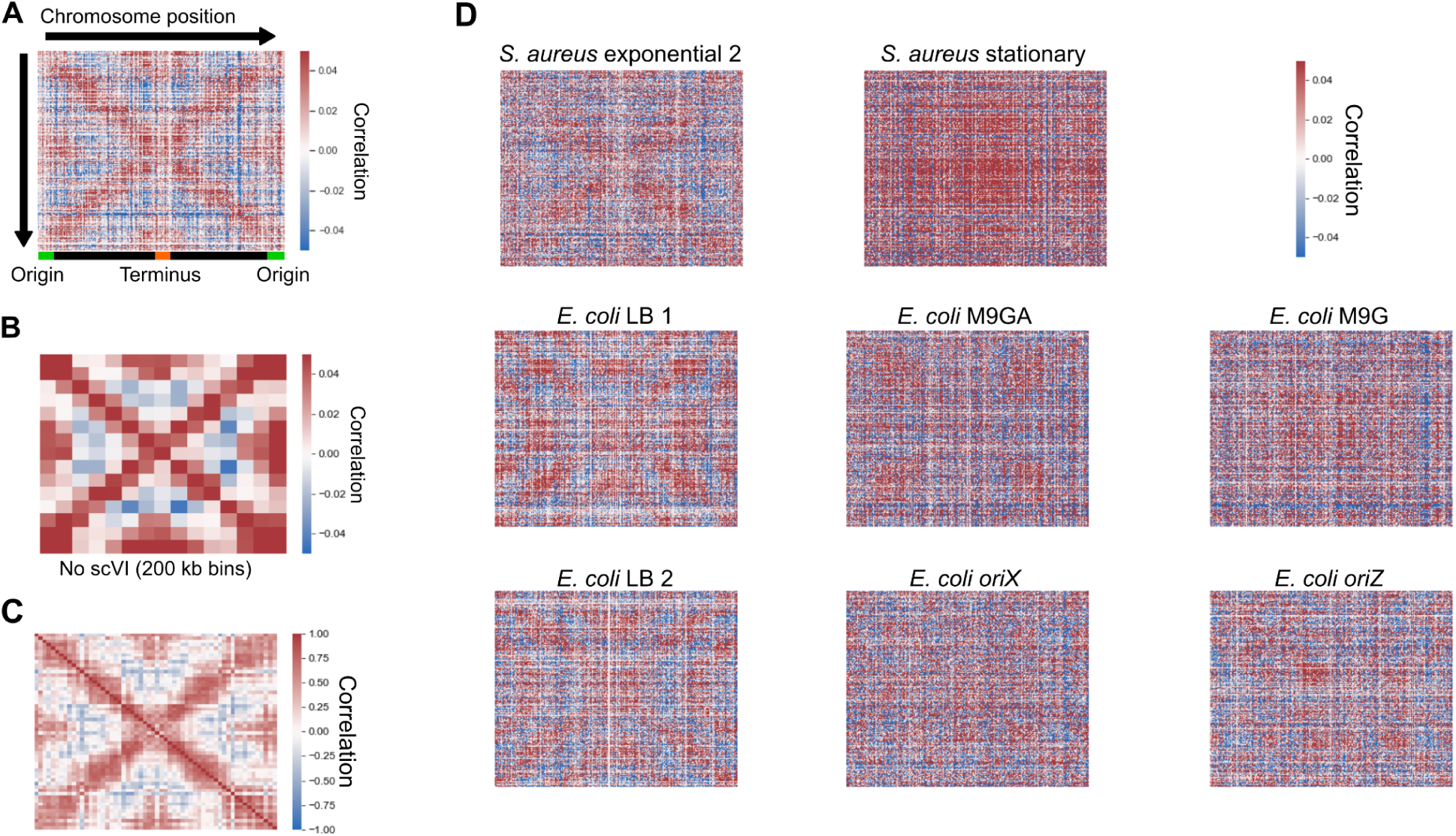
Chromosome-wide gene-gene correlation patterns. **A)** Spearman correlations from Fig. 1C without binning by chromosome position. **B)** Correlations from Fig. 1C without the use of scVI, binning in 200 kb bins by chromosome position. **C)** Spearman correlations in exponential *S. aureus* data from Dataset D4, averaged in 50 kb bins, as for Dataset D3 in Fig. 1C. **D)** Initial correlations from unbinned, scVI-predicted gene expression data. Sample “*S. aureus* exponential 2” is from Dataset D4, whereas *E. coli* LB replicates 1 and 2 are from Dataset D1 and Dataset D2, respectively.

**Figure S3:**
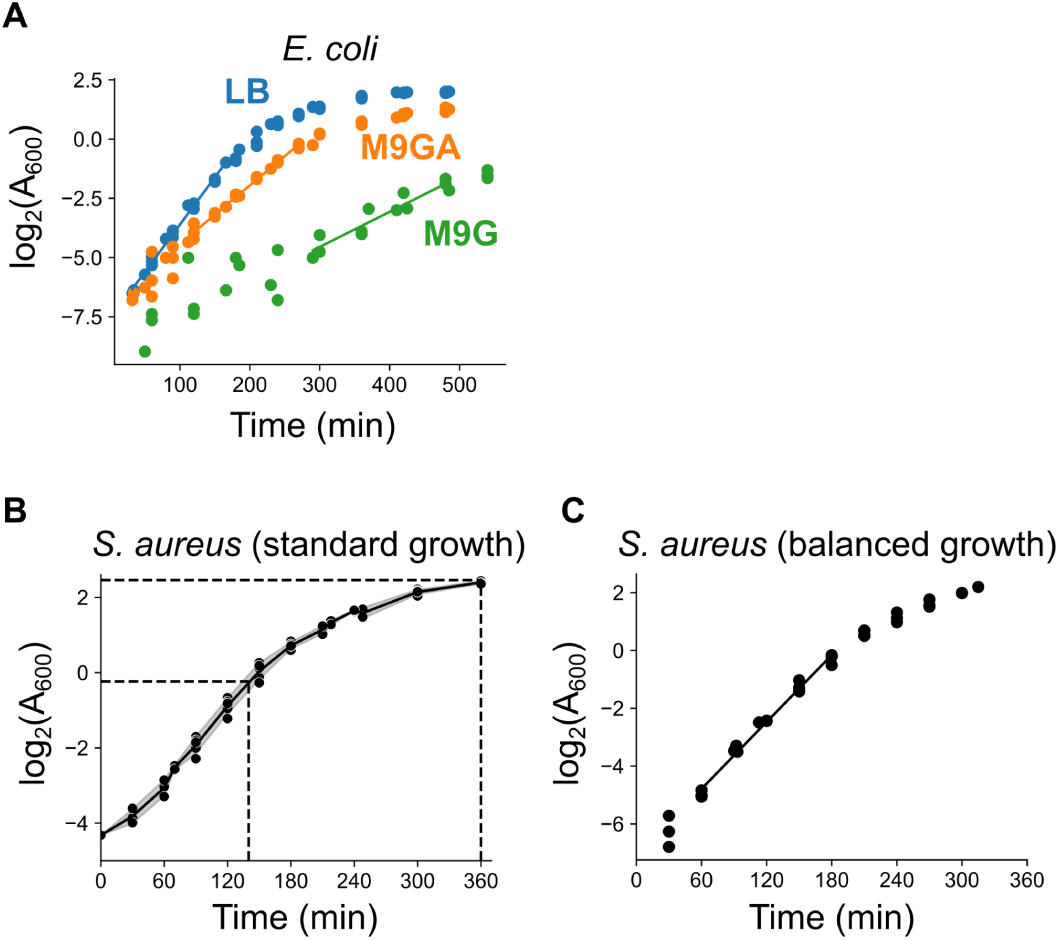
Growth curves of bacterial strains. **A)** Growth of *E. coli* in three conditions. Doubling times were calculated based on the linear portions of growth (marked as fitted lines). Data are from four (LB and M9GA) or three (M9G) biological replicates. **B)** Growth of *S. aureus* under standard growth conditions. The time and log_2_(A_600_) values when exponential and stationary phase samples were taken are marked with dotted lines. The line is fitted to the mean at each time point, with the gray area representing standard deviation. Data are from five biological replicates. Doubling times for exponentially growing cells are estimated for the linear portion of the curve (∼60-150 min). **C)** Growth of *S. aureus* under balanced growth conditions (see Materials & Methods). The black line indicates the linear portion from which doubling time was estimated. Data are from three biological replicates.

**Figure S4:**
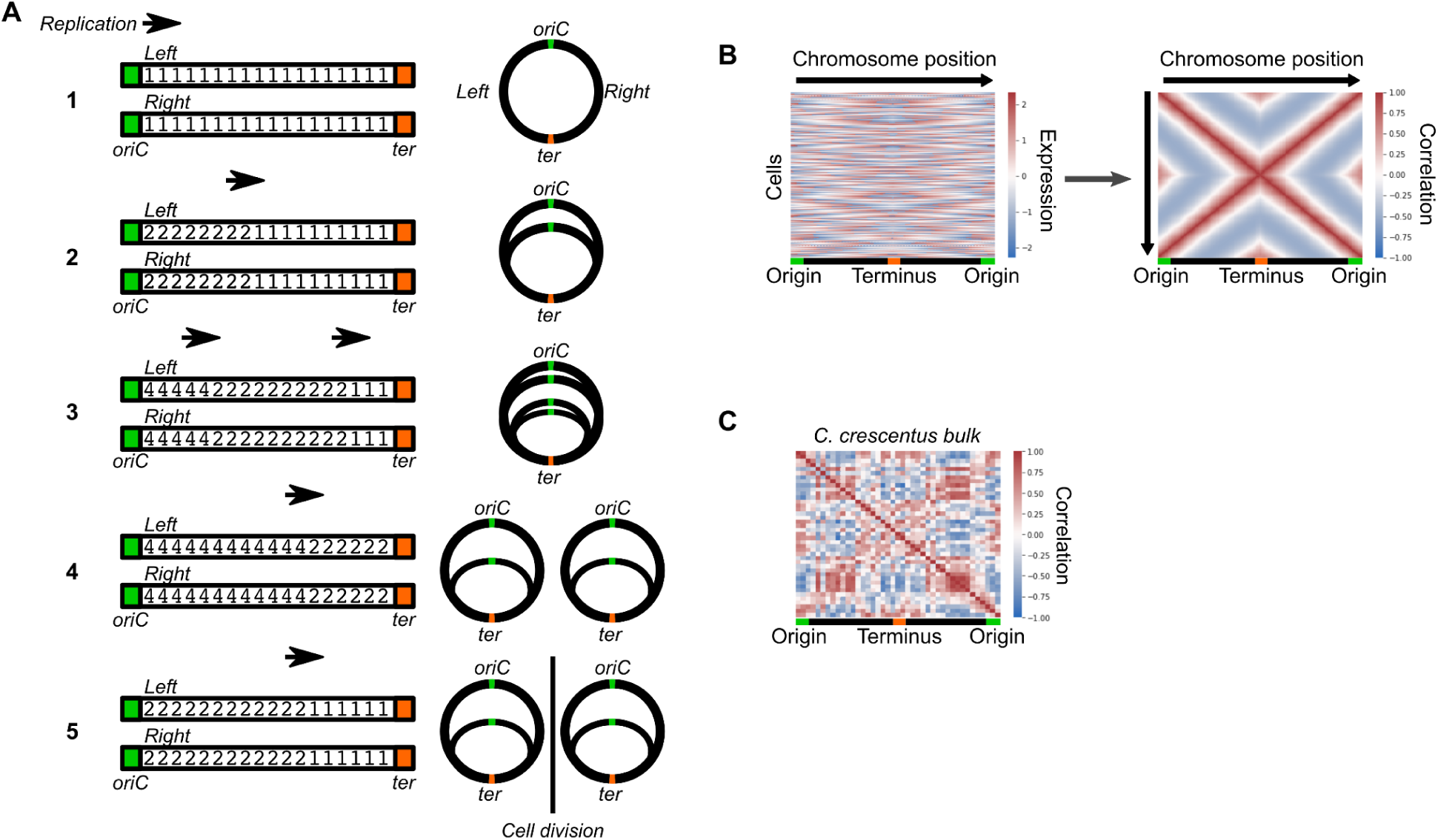
Simulation of replication-dependent gene-gene correlation patterns. **A)** Schematic figure of the simulation. Each “arm” of the circular chromosome is represented as an array of integers (initially ones), representing each gene. Replication proceeds stepwise from origin to terminus, doubling copy number as it does (steps 1 to 2). At high replication rates, a second round of replication will initiate before the first has finished (step 3). When one round of replication reaches the terminus, that round finishes and after a given time interval copy numbers are globally halved, reflecting cell division (steps 4 to 5). Figures on the right indicate the represented states on the circular chromosome. See Materials & Methods for details. **B)** Simulation of DNA copy number effects predicts the global gene covariance pattern. For 1,000 simulated, unsynchronized cells where the doubling time *t_d_* is equal to the C-period, the normalized, scaled gene expression matrix (*left*) is used to calculate gene-gene correlations (*right*). **C)** Gene expression correlations in synchronized *C. crescentus* bulk RNA-seq from ^11^. Scaled gene expression is averaged into 100 kb bins.

**Figure S5:**
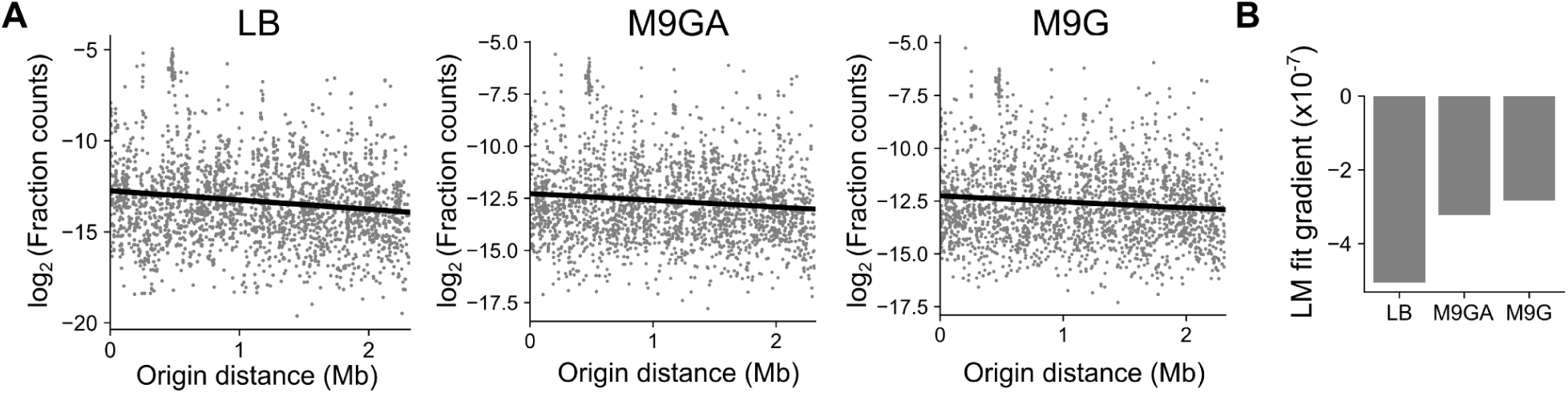
The relationship between origin distance and expression levels. **A)** For each *E. coli* growth condition, the average fraction of total mRNA UMI from each gene was calculated and log_2_-transformed. A linear regression model (black line) was fitted between log-fraction counts and origin distance. **B)** The gradient of the linear model fits in **(A)**. Note that in each case, there is a negative relationship, with a steeper gradient for faster growth rates. This is expected given that at fast growth rates, genes near the origin may attain higher copy number states (>2) than at slow growth rates. Spearman correlations are −0.13 (LB, *P* = 3.8 x 10^-10^), −0.09 (M9GA, *P* = 2.2 x 10^-5^), and −0.07 (M9G, *P* = 6.0 x 10^-4^).

**Figure S6:**
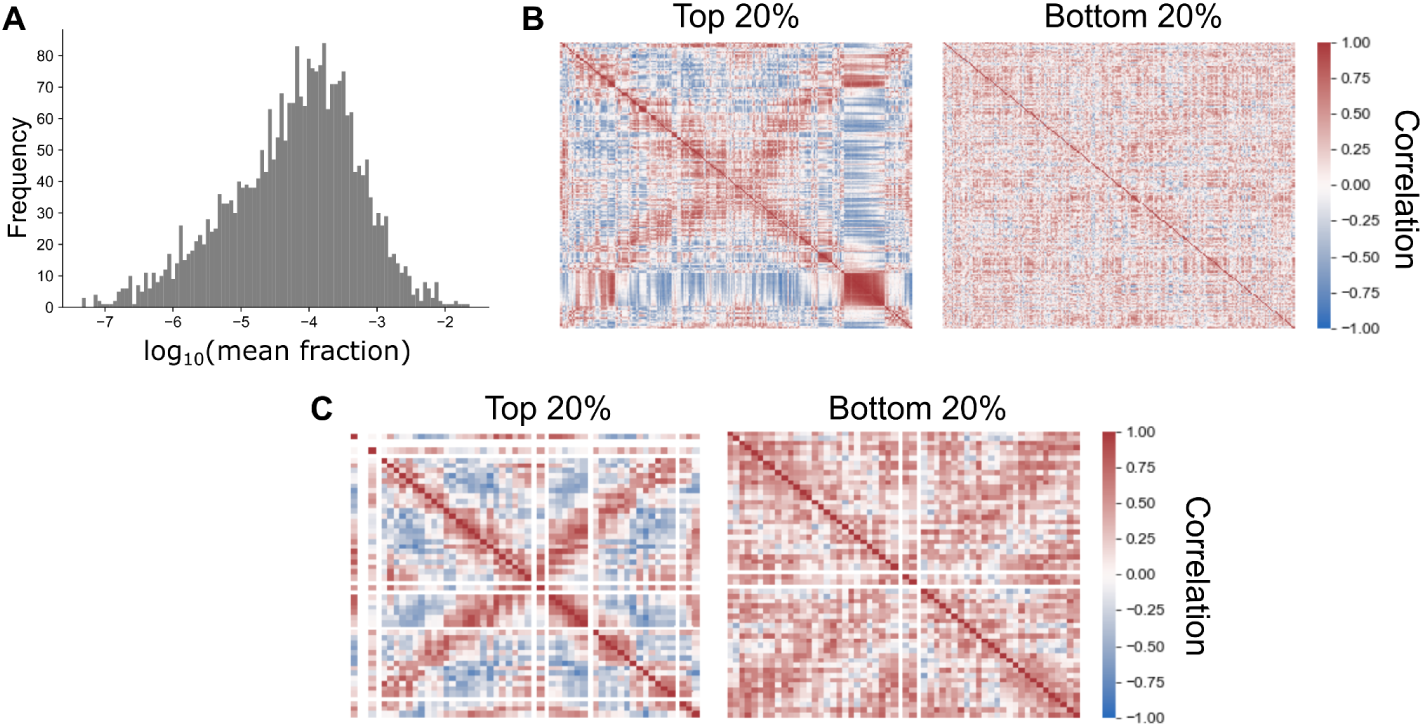
Evidence indicating that the global gene covariance pattern results directly from gene expression. **A)** Histogram showing that length-adjusted average gene expression varies over several orders of magnitude. This is a broad distribution that would not be expected from genomic DNA. Raw expression counts were normalized by library size (to sum to 1 per barcode) and the average expression was calculated. Length correction was performed as expression divided by gene length then multiplied by median gene length. **B)** Spearman correlations between genes in the top and bottom 20% of genes. Genes are arranged by chromosome order. **C)** Spearman correlations between top and bottom 20% of genes after averaging expression in 50 kb bins as in Fig. 1C. For **(C & D)**, if the pattern was driven by low-level contaminating genomic DNA, it would be expected to be more evident in low-expressed genes (since a higher proportion of reads from these genes should come from genomic DNA) than in high-expressed genes. The opposite is true, with a much stronger pattern in high-expressed genes (presumably due to less noise in these measurements). Taken together, these observations strongly support that the pattern is driven by variation in the transcriptome rather than contaminating genomic DNA.

**Figure S7:**
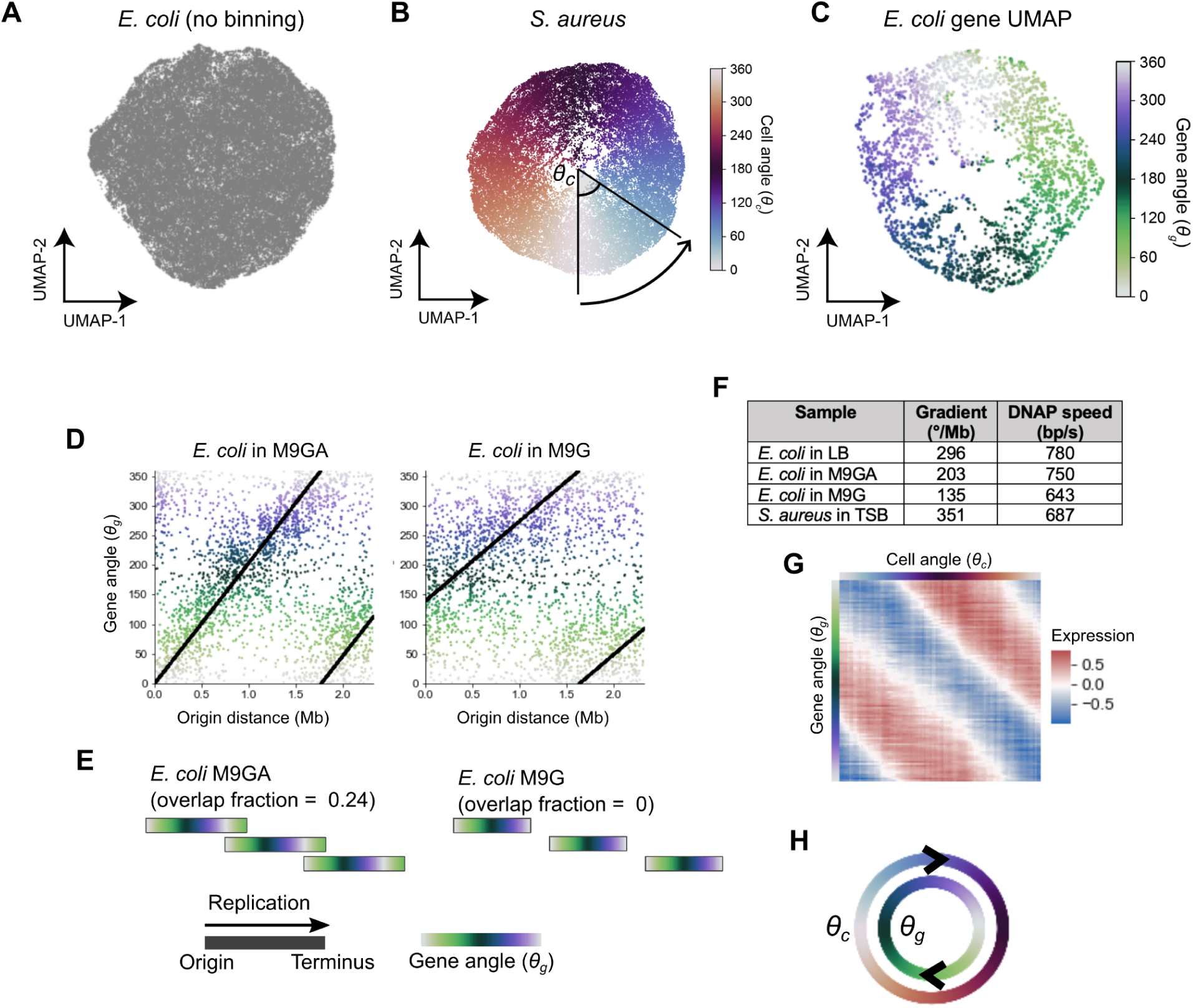
Cell and gene angle analysis to model replication-dependent gene expression. **A)** UMAP analysis of LB-grown *E. coli* based on scVI-predicted expression. **B)** UMAP of *S. aureus* with gene expression averaged in 50 kb bins by chromosome position. Cells are colored by the cell angle *θ_c_* between UMAP dimensions relative to the center of the projection. **C)** UMAP of *E. coli* genes, performed on the same data as the PCA in Fig. 2D. Gene angles shown are those derived from PCA. **D)** The relationship between *θ_g_* and origin distance for *E. coli* grown in M9 + glucose + amino acids (M9GA) or M9 + glucose (M9G). The black line indicates the model fit as described in Materials & Methods Section “Modeling the gene angle-origin distance relationship”. **E)** Predicted replication patterns as for Fig. 2G but for *E. coli* under slower growth conditions. **F)** Gradients of the gene angle-origin distance relationship and estimates of DNA polymerase speed from these gradients. See Materials & Methods for details. **G)** Expression in LB-grown *E. coli* is first averaged in 100 bins by *θ_c_* then averaged in 100 bins by *θ_g_* to yield the 100 x 100 matrix represented here as a heatmap. This is used to train the model to predict gene expression at a given point in the cell cycle (*θ_c_*) for a given gene (*θ_g_*). **H)** Conceptual representation of the cell cycle expression parameterization. Cells are ordered in their cell cycle state by *θ_c_*, whereas genes are ordered by their cell cycle expression by *θ_g_*. Cell cycle expression can be described as the concurrent cycling of cells and genes ordered by these metrics.

**Figure S8:**
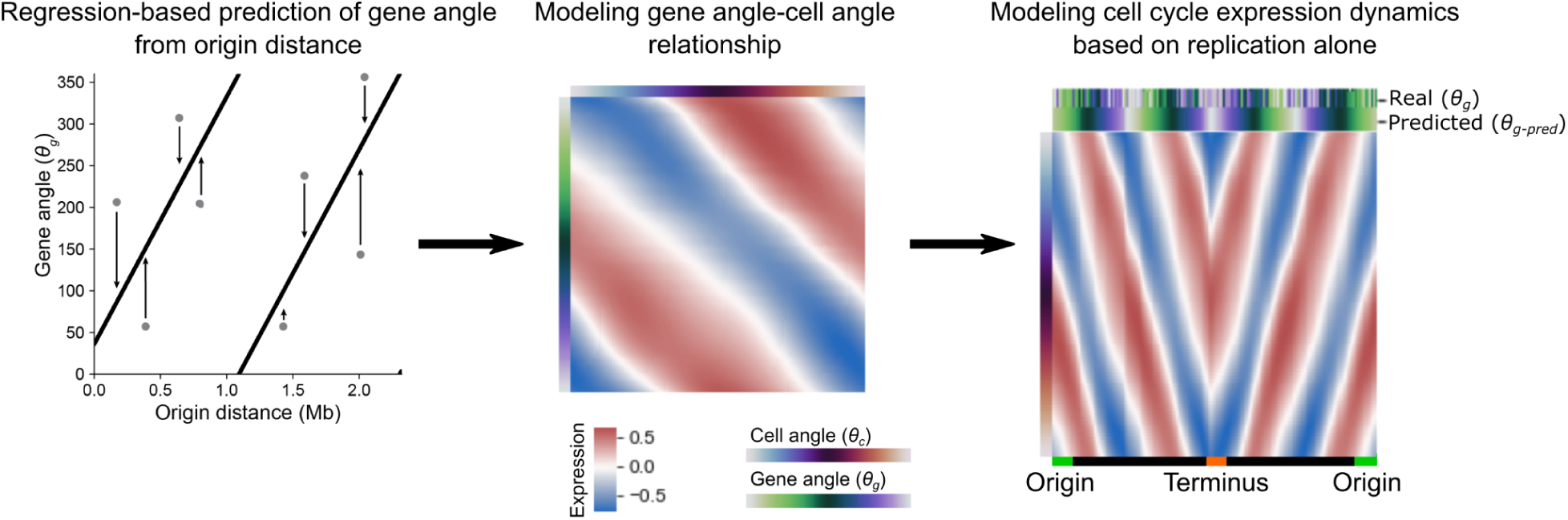
Predicting gene expression dynamics based on distance from the origin. The following pipeline predicts cell cycle expression for a given gene based only on its distance from the origin of replication. A regression model predicts gene angle *θ_g-pred_* based on origin distance alone (*left*) and this is converted into a prediction of expression by cell angle *θ_c_* using a second regression model (*middle*). Ordering genes by chromosome position (*right*) shows a smoothed version of the expression pattern in Fig. 2B. The bar at the top of this figure shows the real and predicted gene angles. Data are from *E. coli* grown in LB. See Materials & Methods for full details.

**Figure S9:**
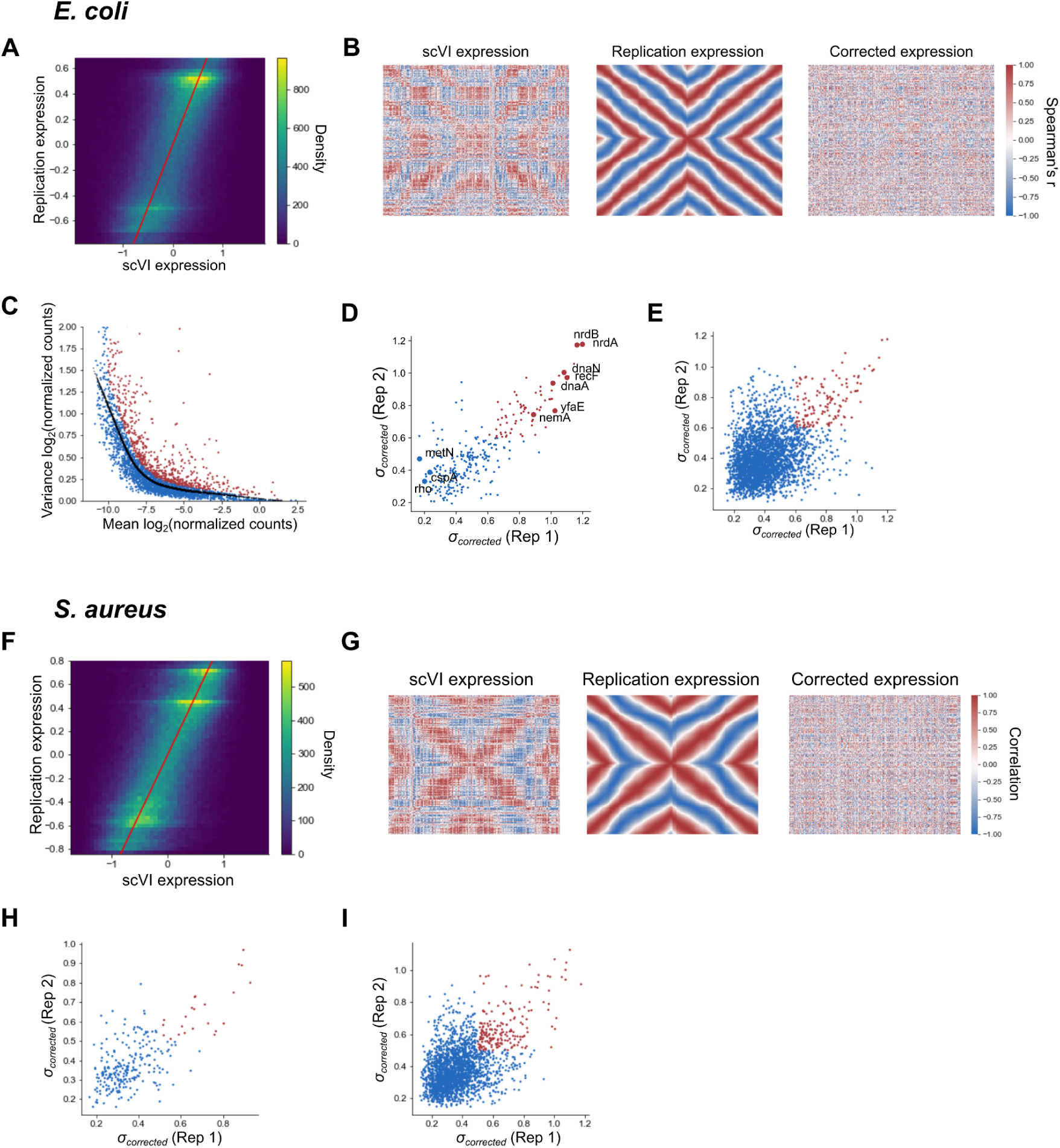
Correcting for and measuring divergence from predicted replication-associated patterns. **A)** Two-dimensional histogram for *E. coli* showing the relationship between observed expression from scVI and replication-predicted expression. Expression is averaged in 100 bins by cell angle *θ_c_*. The red line indicates *x* = *y* i.e. the case where expression in both matrices is identical. Overall, there is a rough 1:1 correspondence between observed and predicted expression, indicating a good model fit. **B)** Gene-gene correlations in LB-grown *E. coli* across *θ_c_*-binned expression data (100 bins) for the full scVI observed model (*left*), the replication-only model (*middle*), and the corrected model that is the difference of the two expression matrices (*right*). **C)** The mean-variance relationship in *E. coli* of log-transformed normalized counts. The black line indicates the locally weighted scatterplot smoothing (LOWESS)-fitted values and red points are genes classed as highly variable. See Materials & Methods for further details. **D)** Comparison of the divergence score *σ_corrected_* between LB-grown *E. coli* in Datasets D1 & D2 of genes classed as highly variable in both datasets (287 genes). Red indicates replication-divergent genes (*σ_corrected_* > 0.6). **E)** Comparison of *σ_corrected_* (standard deviation of divergence from the replication model) between LB-grown *E. coli* in Dataset D1 and Dataset D2 of all genes present in both datasets. Red indicates *σ_corrected_* > 0.6 in both datasets, meaning that they are considered replication-divergent. The Pearson correlation between replicates is 0.38. **F)** Two-dimensional histogram as in **(A)** but for *S. aureus*. **G)** Gene-gene correlation plots as for **(B)** but for *S. aureus*. **H & I)** Comparison of *σ_corrected_* (standard deviation of divergence from the replication model) between *S. aureus* in Dataset D5 and Dataset D6 for highly variable genes in both datasets **(H)** (Pearson’s *r* = 0.66) and all genes **(I)** (Pearson’s *r* = 0.48). Red indicates *σ_corrected_*> 0.5 in both datasets, meaning that they are considered replication-divergent.

**Figure S10:**
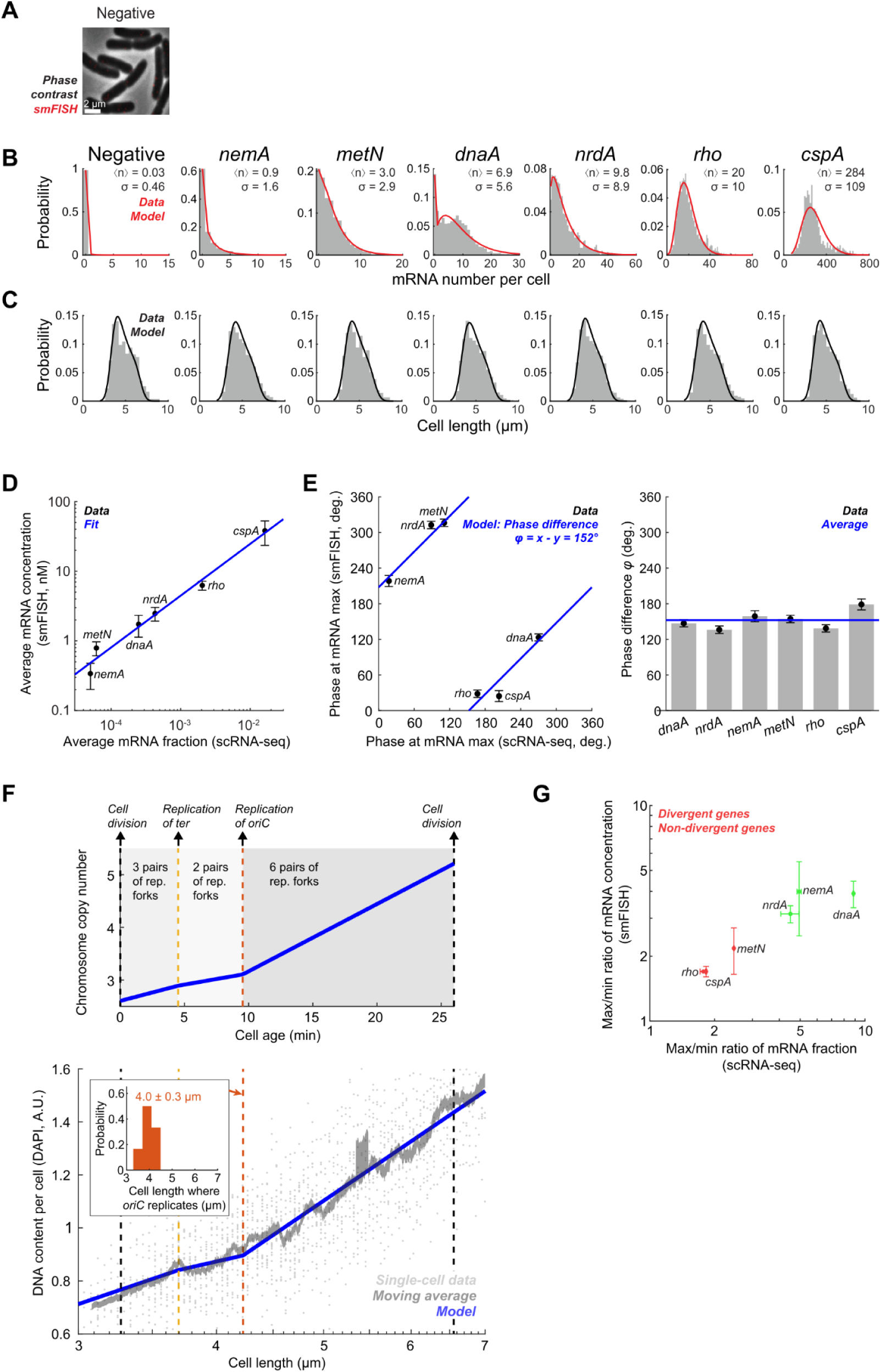
smFISH analysis of cell cycle gene expression correlates with phase-shifted scRNA-seq data. **A)** Negative control for smFISH labeling. *E. coli* cells labeled against bacteriophage lambda *cI* mRNA. smFISH signal is shown using the same contrast as in Fig. 3, B & D. See Section “smFISH”. **B)** The distribution of mRNA copy-number per cell for each gene. See Section “mRNA quantification”. Red line, fit to a negative binomial distribution plus a “zero spike”^6^. **C)** The distribution of cell length in each sample. Black line, fit to the theoretical model of ^84^, see Section “Modeling the distribution of cell length”. **D)** Comparison of the population-averaged mRNA fraction, as measured using scRNA-seq, with mRNA concentration, as measured using smFISH. Markers and error bars indicate mean ± SD from two datasets of each method. Blue line, fit to a function y = ax^b^. **E)** Estimation of the cell-cycle phase difference between scRNA-seq and smFISH. The phase of each dataset was estimated as described in Section “Cell-cycle analysis of mRNA concentration”. *Left*, markers and error bars indicate mean ± SEM from two datasets of each method. Blue line, fit to a linear function, indicating a constant phase difference φ. *Right*, the estimated phase difference across the six genes examined. **F)** Top, the theoretically predicted cellular DNA contents as a function of cell age, see Section “Inferring cell-cycle phase from the DAPI signal”. Bottom, DAPI-measured DNA content per cell as a function of cell length. Single-cell data was binned based on cell length (moving average ± SEM, 21 cells per bin), Blue line, fit to the theoretical model. Inset, the distribution of the inferred cell length where *oriC* replicates, estimated from all smFISH samples. **G)** Divergent genes exhibit a larger amplitude of cell-cycle fluctuations. The ratio between the maximum and minimum expression level of different genes, as measured using scRNA-seq and smFISH. The mRNA fraction (scRNA-seq) and concentration (smFISH) were obtained as in Fig. 3 B & D, 2nd and 3rd columns. The maximum and minimum levels were determined from the binned data. Markers and error bars indicate mean ± SD from two datasets of each method.

**Figure S11:**
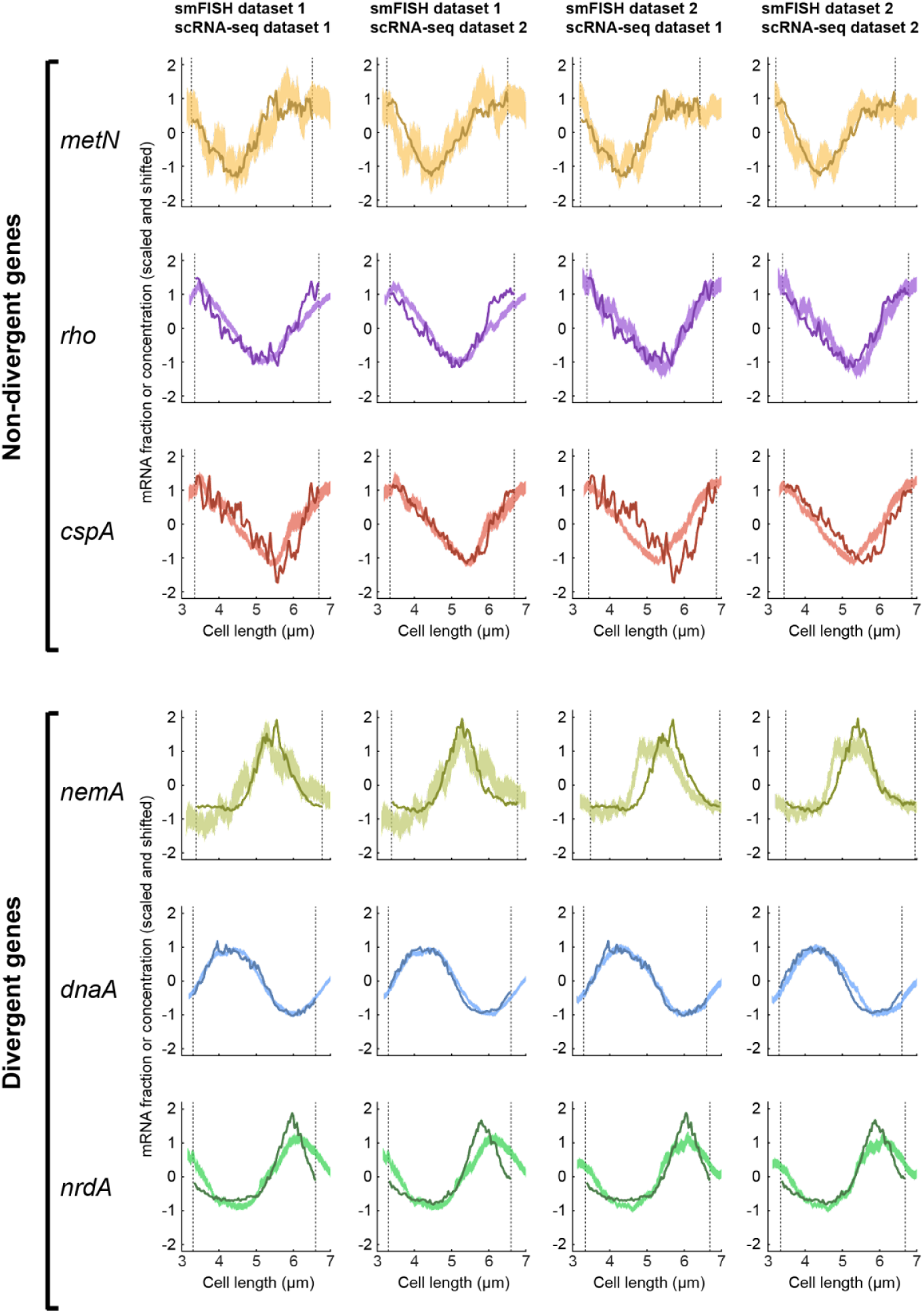
Cell cycle analysis of smFISH and scRNA-seq shows good agreement across biological replicates. Pairwise comparison between two smFISH and two scRNA-seq datasets. Analysis as in Fig. 3B & D, 4th column. See Section “Cell-cycle analysis of mRNA concentration”.

**Figure S12:**
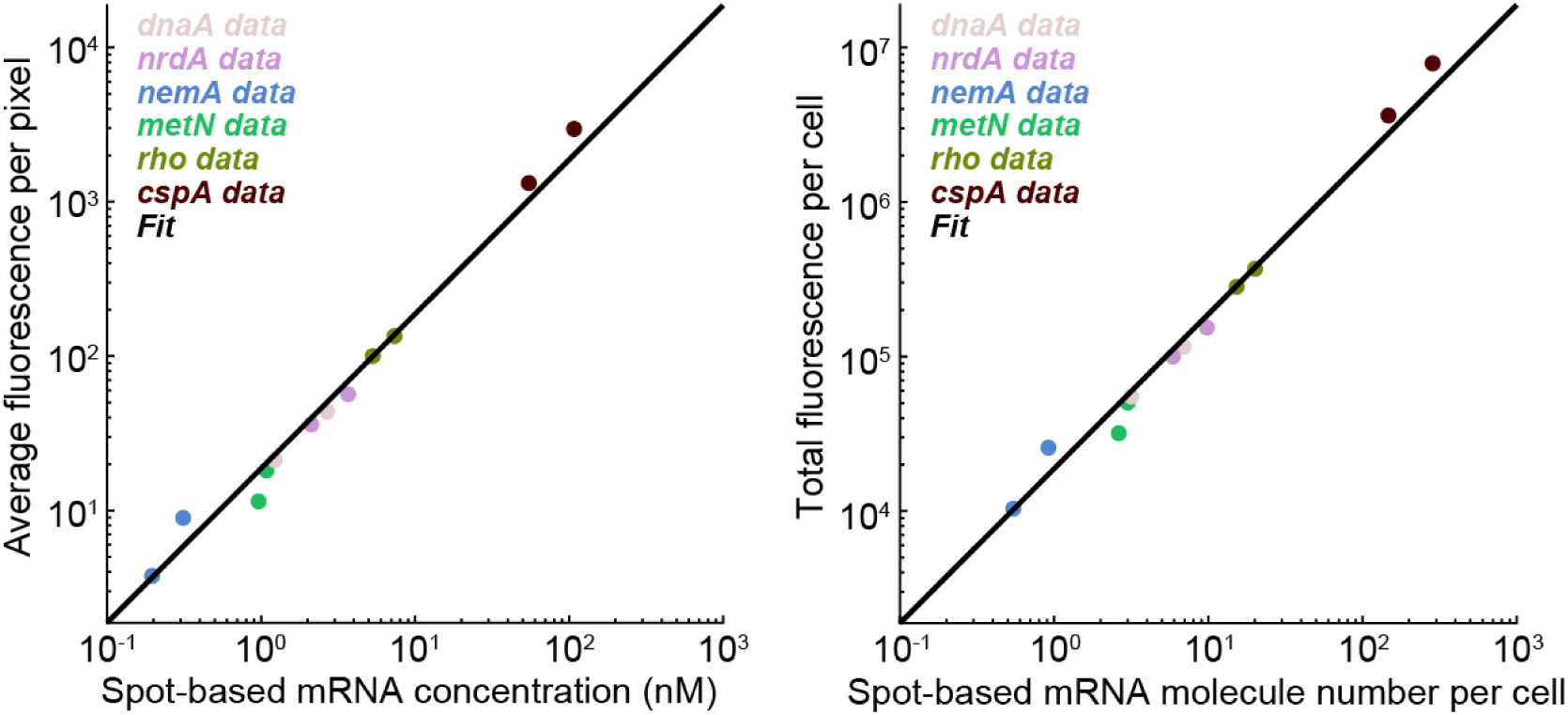
Consistency between spot-based and cell-based smFISH quantification. Comparison of the mRNA levels inferred from smFISH data using spot-based and cell-based mRNA quantification. Both methods are described in Section “mRNA quantification”. *Left*, mRNA concentration. *Right*, mRNA copy number per cell. Markers indicate mean values from each smFISH sample (Error bars are smaller than marker size). Black line, fit to a linear function.

**Figure S13:**
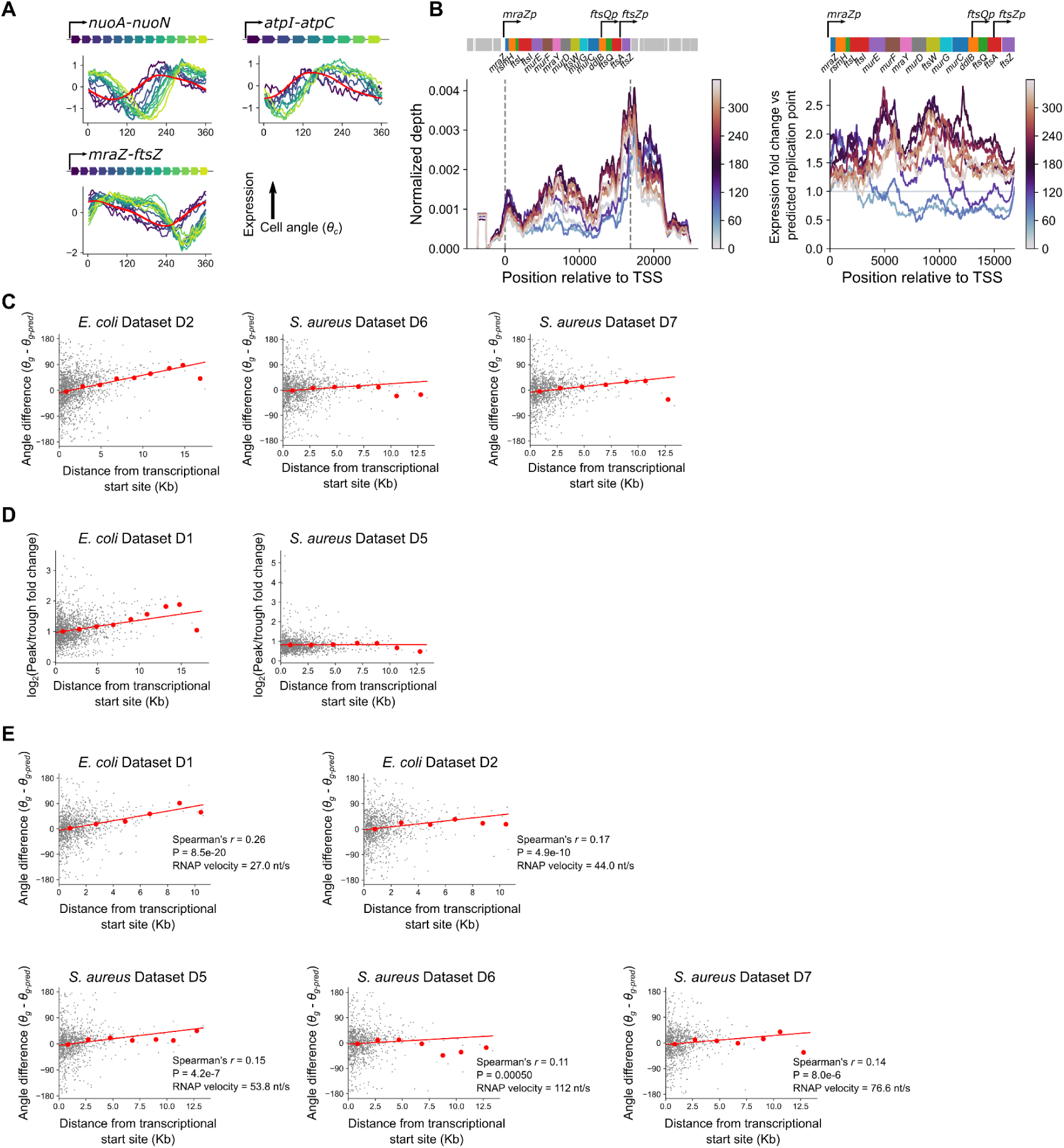
The relationship between distance from the transcriptional start site and gene expression timing and amplitude. **A)** Cell cycle gene expresion plots for operons showing “delayed” genes as in Fig. 4B but for LB-grown WT *E. coli* from Dataset D2. The red line indicates predicted expression. **B)** Normalized per-base read depth at the *mraZ-ftsZ* locus. *Left*: Normalized expression as in Fig. 4D. *Right*: Fold-change relative to expression at the predicted time of replication, as in Fig. 4E. Schematic figures of the locus depict a simplified version since several internal promoters have been identified. **C)** Plots of maximum distance from a transcriptional start site against difference between predicted and observed angles as in Fig. 4C. Red line indicates the linear model fit and red points indicate averages of 2 kb bins. Data are shown for additional *E. coli* and *S. aureus* replicates. **D)** Plots as in **(C)** but of maximum distance from a transcriptional start site against the log_2_-transformed peak/trough ratio in gene expression, calculated as described in Materials & Methods. **E)** Plots as in **(C)** but using manual operon annotation. Here, any tandem, contiguous stretch of genes with an intergenic distance less than 40 bp is considered an operon. Transcriptional start sites are defined as the start position of the first gene in the operon.

**Figure S14:**
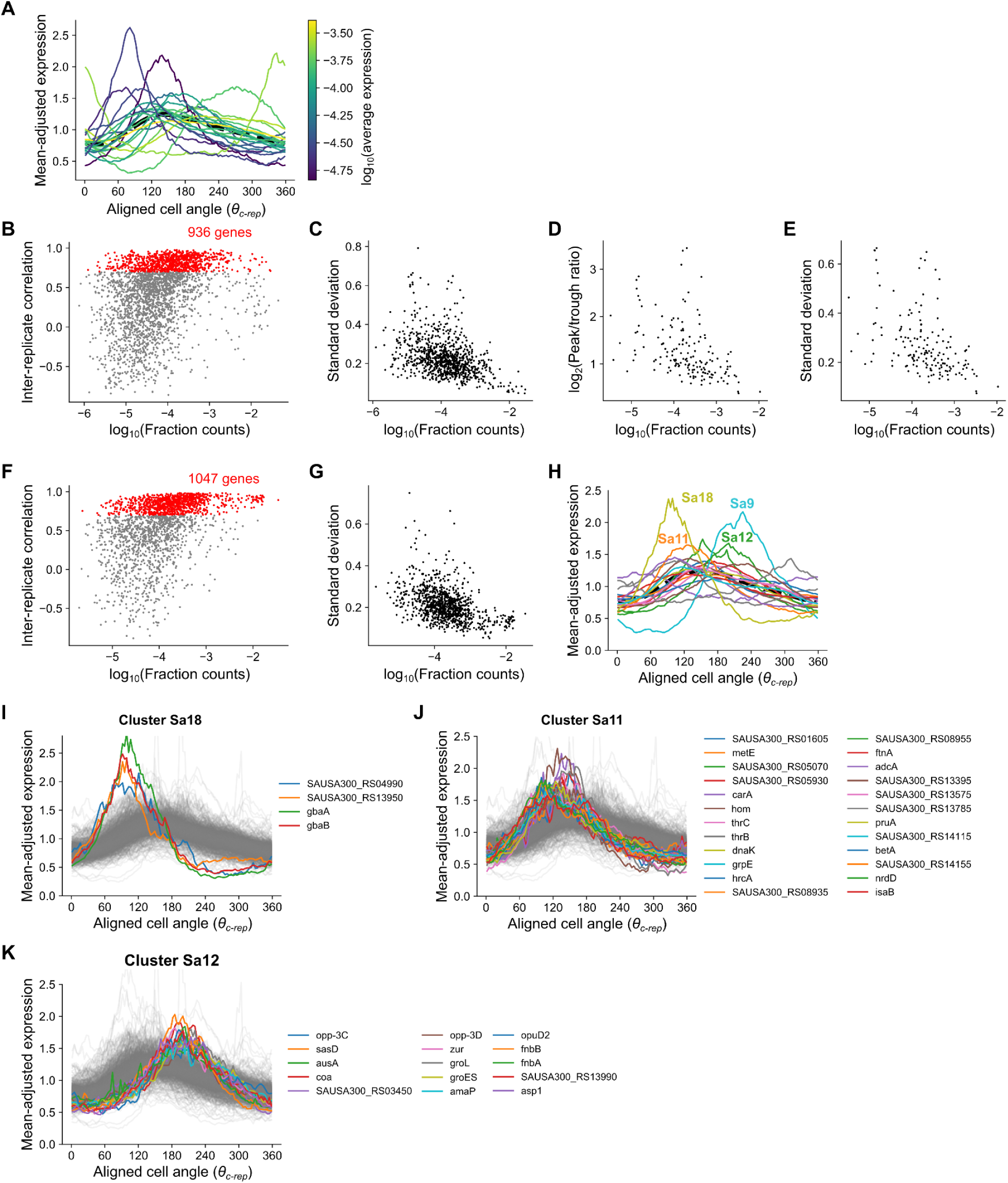
Expression-amplitude relationships and *S. aureus* cluster profiles. **A)** Clusters as in Fig. 5B but colored according to their average, length-corrected expression. This was determined by a gene’s mean fraction of total mRNA that was length-corrected by dividing by its length and multiplying by the median gene length across genes. **B)** Scatter plot of length-corrected mean fraction counts (i.e. fraction of a gene within the whole transcriptome) against Spearman correlation in *E. coli*. Spearman correlations for each gene were calculated as the inter-replicate correlation between cell cycle gene expression measurements averaged in 100 bins by *θ_c_* (replicates from Datasets D1 & D2). Red genes indicate the reproducible genes used in Fig. 5. **C)** Length-corrected mean expression against standard deviation across expression averaged in 100 bins by *θ_c_*. **D & E)** Plots as in Fig. 5E and **(C)** but including only those genes with Spearman R > 0.9 (instead of 0.7). **F)** Plot as in **(B)** but for *S. aureus* (replicates from Datasets D5 & D6). **G)** Plot as in **(C)** but for *S. aureus*. **H)** Plot as in Fig. 5B except for mean expression of *S. aureus* clusters. Genes situated on mobile genetic elements were removed prior to clustering analysis. **I-K)** Plots of individual genes from clusters indicated in **(H)**.

**Figure S15:**
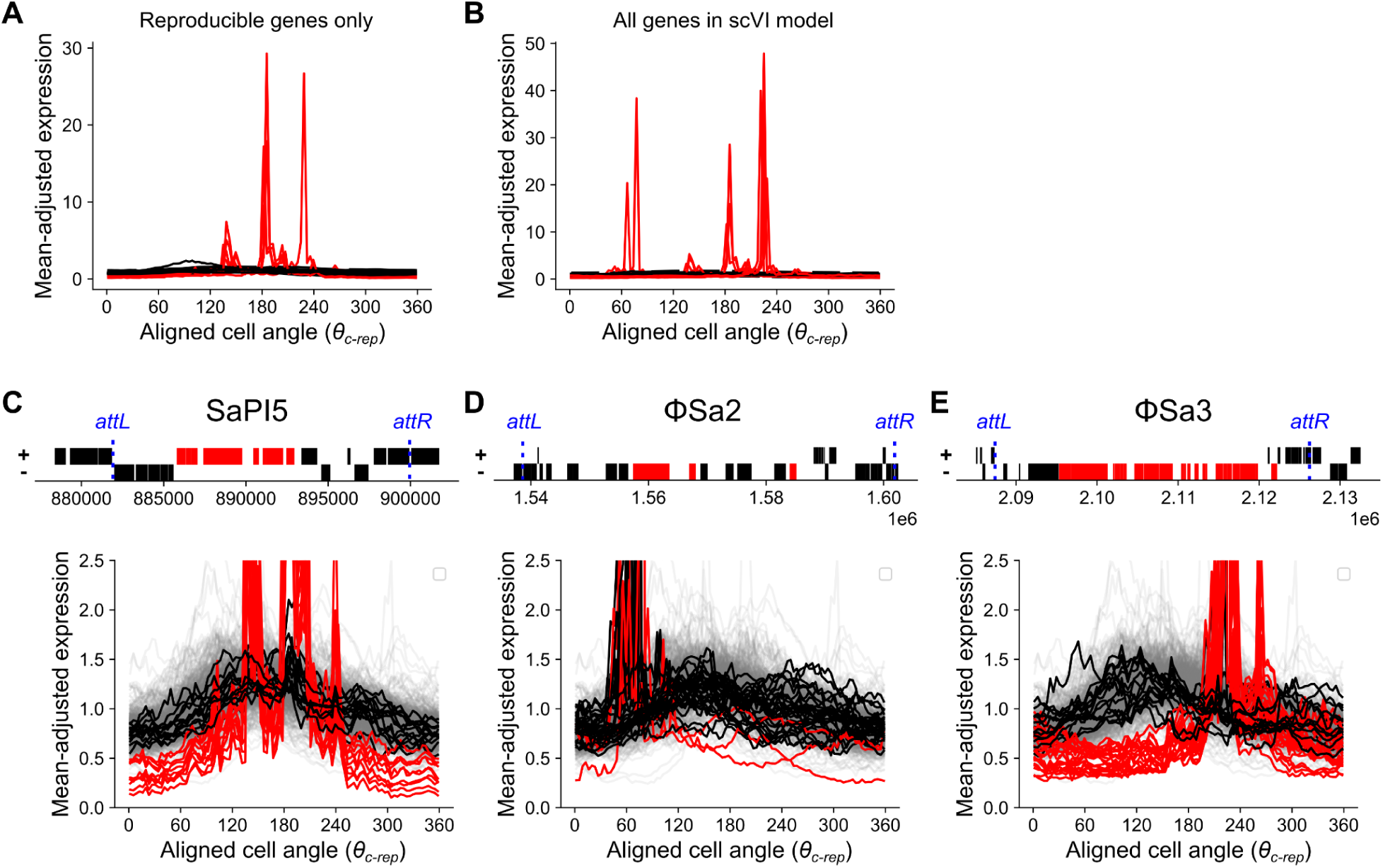
Core genes of mobile genetic elements show highly divergent expression patterns. **A)** Mean cluster expression of reproducible genes partitioned into 20 clusters by aligned gene expression (*θ_c-rep_*). Clusters in red are those that only contain genes located within MGEs. **B)** Plot as in **(A)** but with clustering performed on all genes included in the scVI model (regardless of reproducibility). These cluster assignments are used for **(C-E)**. **C-E)** Expression of genes within mobile elements. Genes are colored based on whether they are in MGE-exclusive clusters from **(B)** (red) or other clusters (black). *Top*: schematic figure of MGE gene content. The *x*-axis represents chromosomal coordinate and + and - strands are plotted separately by *y*-axis position. Predicted attachment sites *attL* and *attR* denote the predicted boundaries of the MGE and are annotated in blue. Annotation for MGEs was taken from the online tool Phaster^89^. *Bottom*: Plots of MGE genes by aligned gene expression (*θ_c-rep_*) as represented in Fig. 5. Gray genes represent the non-MGE background. Note that phage ΦSa2 is disabled and expression of its MGE-specific (“red”) cluster genes is low (0.002% of cells contain at least three transcripts) compared to the staphylococcal pathogenicity island (SaPI) 5 (0.4%) and phage ΦSa3 (0.07%), potentially contributing to the less clear delineation between expression profiles by gene type. MGE-specific expression patterns may arise due to MGE mobilization and these patterns may represent rare events that are not effectively captured by our cell cycle analysis, meaning that the plots here should be interpreted with caution.

**Figure S16:**
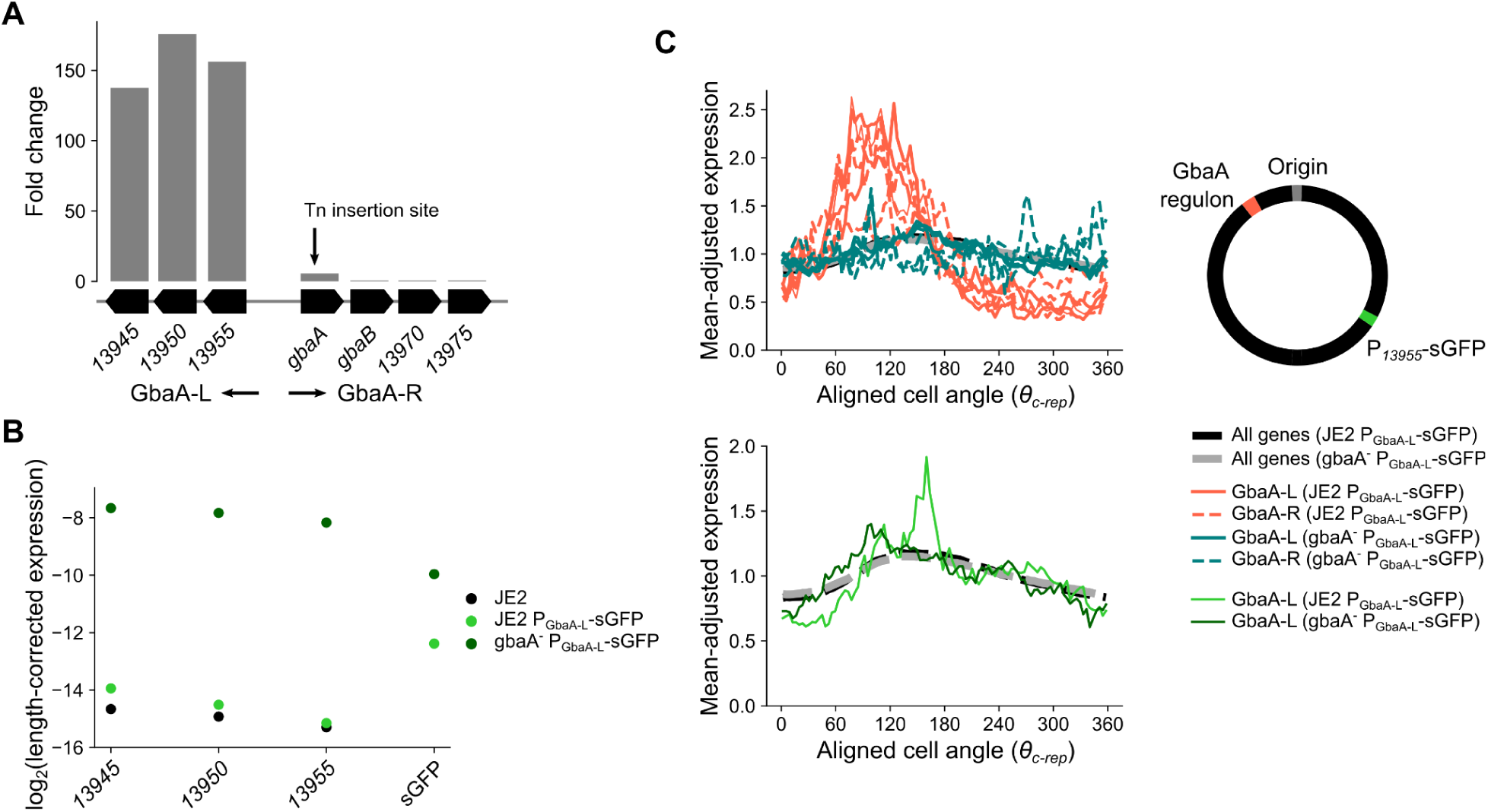
Effects of gbaA disruption on cell cycle gene expression. **A)** Expression fold change of genes in the GbaA regulon after *gbaA* transposon insertion. Genes of the GbaA-L operon increase in expression >100–fold. However, due to the location of the transposon insertion towards the 5’ end of *gbaA*, induction of GbaA-R genes is not observed. Genes with names starting with *SAUSA300_RS* are truncated to give only the unique number. **B)** Average expression of GbaA-L genes and sGFP in reporter strains (compared to JE2 in measurements from the same experiment). Average expression measured as fraction of total mRNA was length-corrected as elsewhere by dividing by the gene length and multiplying by the median gene length across all genes. Note that sGFP expression in JE2 P_GbaA-L_-sGFP is approximately fourfold higher than that of GbaA-L genes, and the derepressed form in *gbaA^-^*P_GbaA-L_-sGFP is also fourfold lower (possibly reflecting lower copy number due to its further distance from the origin). Therefore, while repression of the GbaA-L locus is ∼96-fold, repression of sGFP by GbaA is only 5.3-fold. **C)** Comparison of aligned expression (*θ_c-rep_*) (as in Fig. 5) for GbaA regulon genes and sGFP in the two reporter constructs. Thick black and gray lines represent average expression across all reproducible genes. The schematic figure represents the relative positions of the GbaA regulon and the P_GbaA-L_-sGFP integration site.

**Figure S17:**
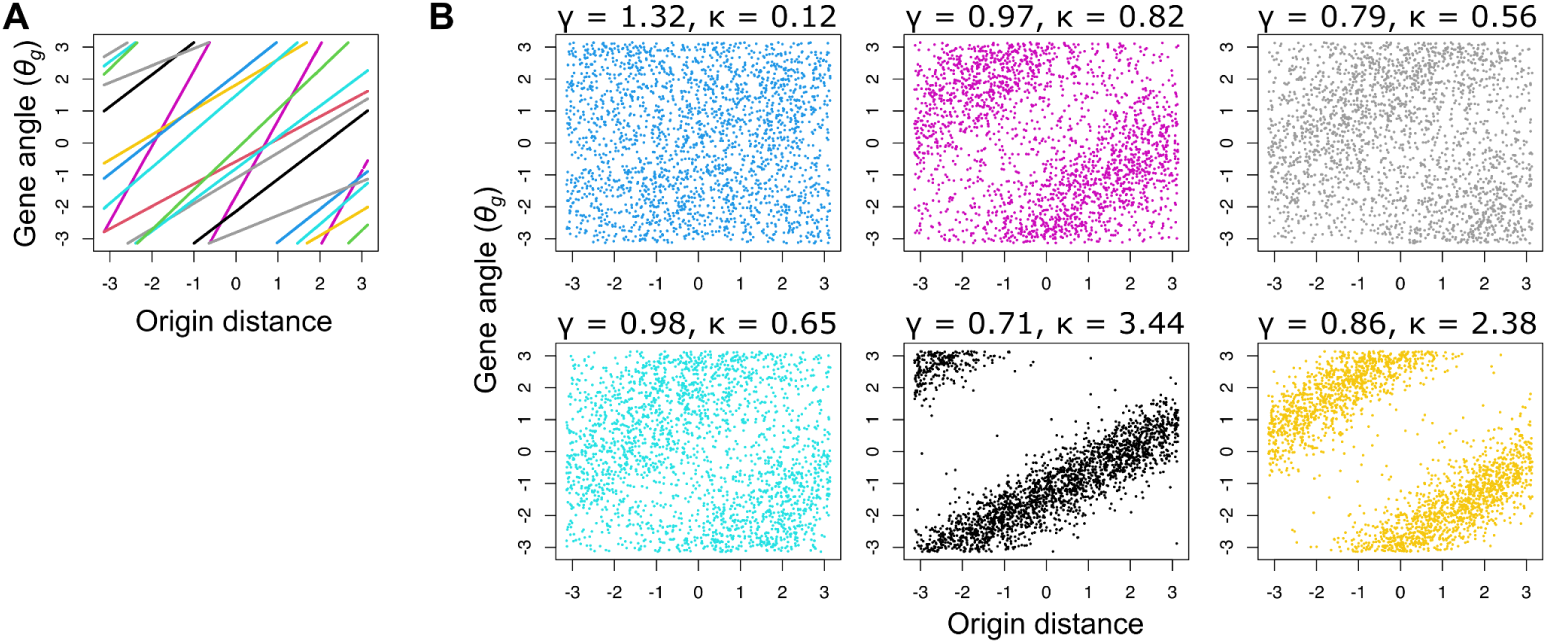
Sampling from the prior of the gene angle-origin distance regression model. Based on the model and priors specified in Materials & Methods, values were randomly sampled from the prior and used to predict either the expected gene angle *A* **(A)** or the predicted value of gene angle *θ_g_* after von Mises sampling **(B)**. For each sampled set of parameters in **(B)** the gradient *ɣ* and concentration parameter *κ* are shown. Both *θ_g_* and origin distance *D* are standardized to the range -π to π as per the model requirements. Overall, the prior assumptions of the model are that there is a positive, linear relationship between *θ_g_* and *D*, but there is considerable flexibility regarding the gradient (and hence degree of wrapping), value of *θ_g_* at *D* = 0, and noise.

**Table S1:**
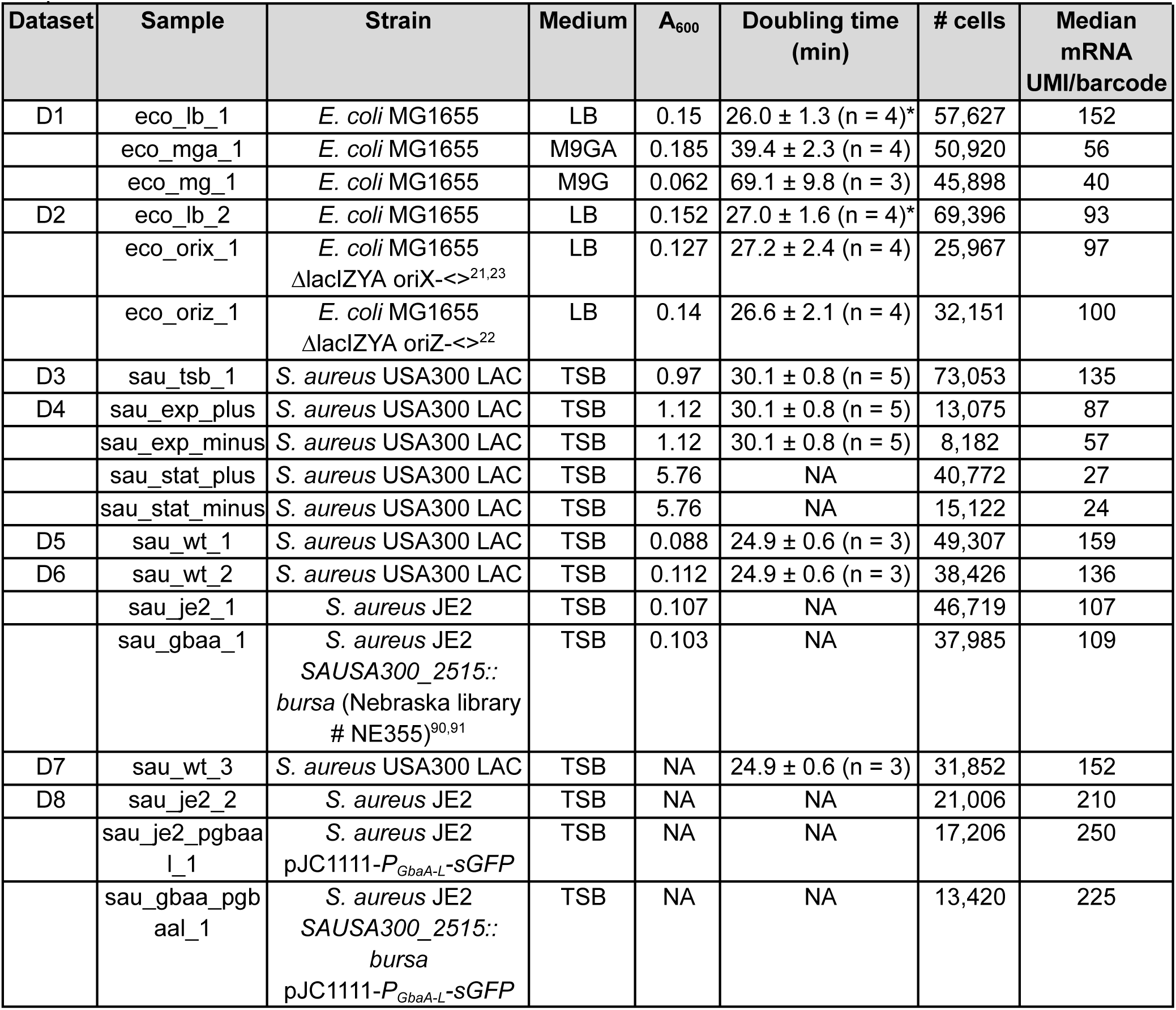
Information about datasets and samples used. A_600_ refers to the optical density at the time of harvesting. *Growth E. coli MG1655 in LB was measured in a separate series of experiments for each dataset.

**Table S2:**
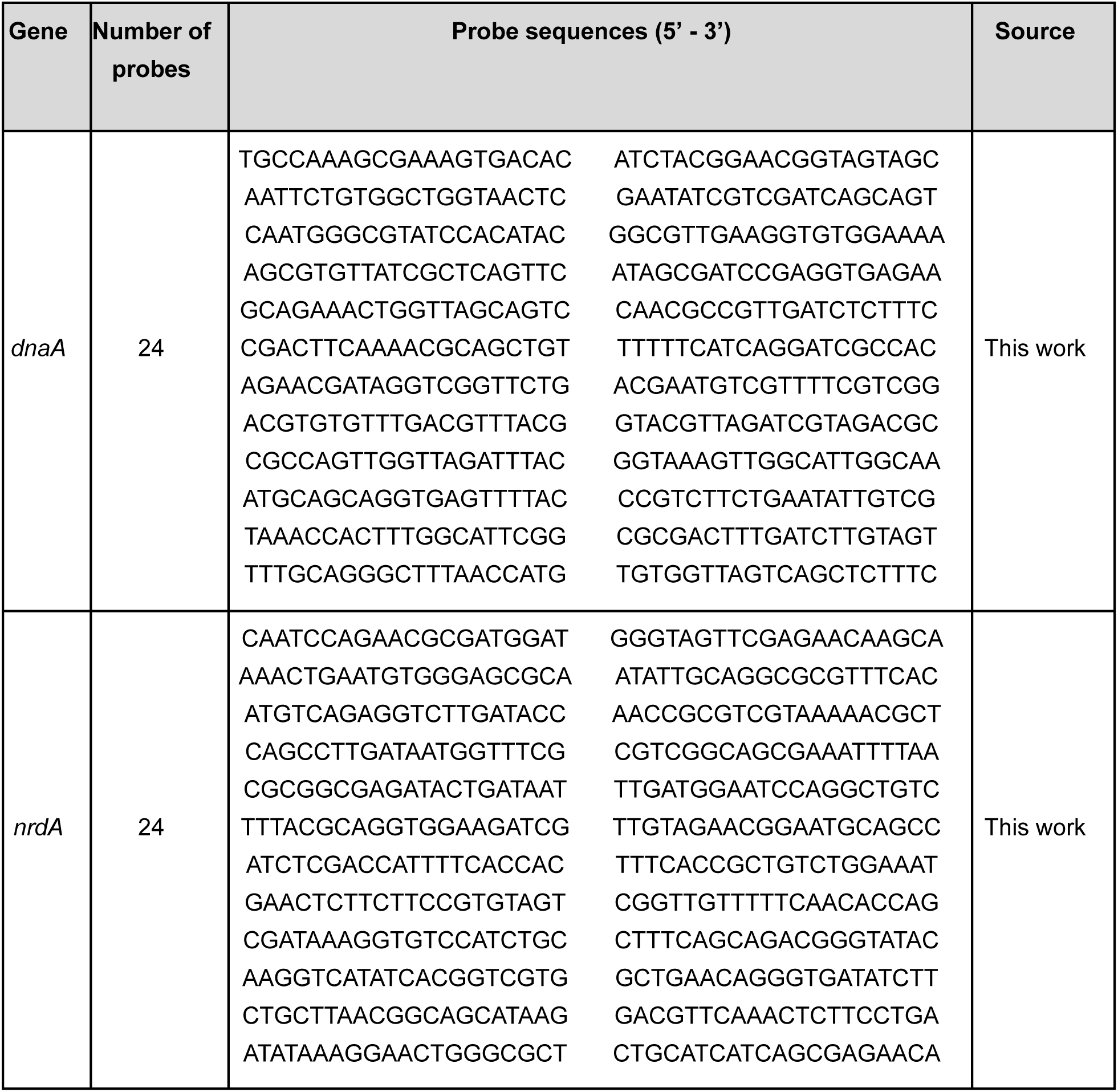
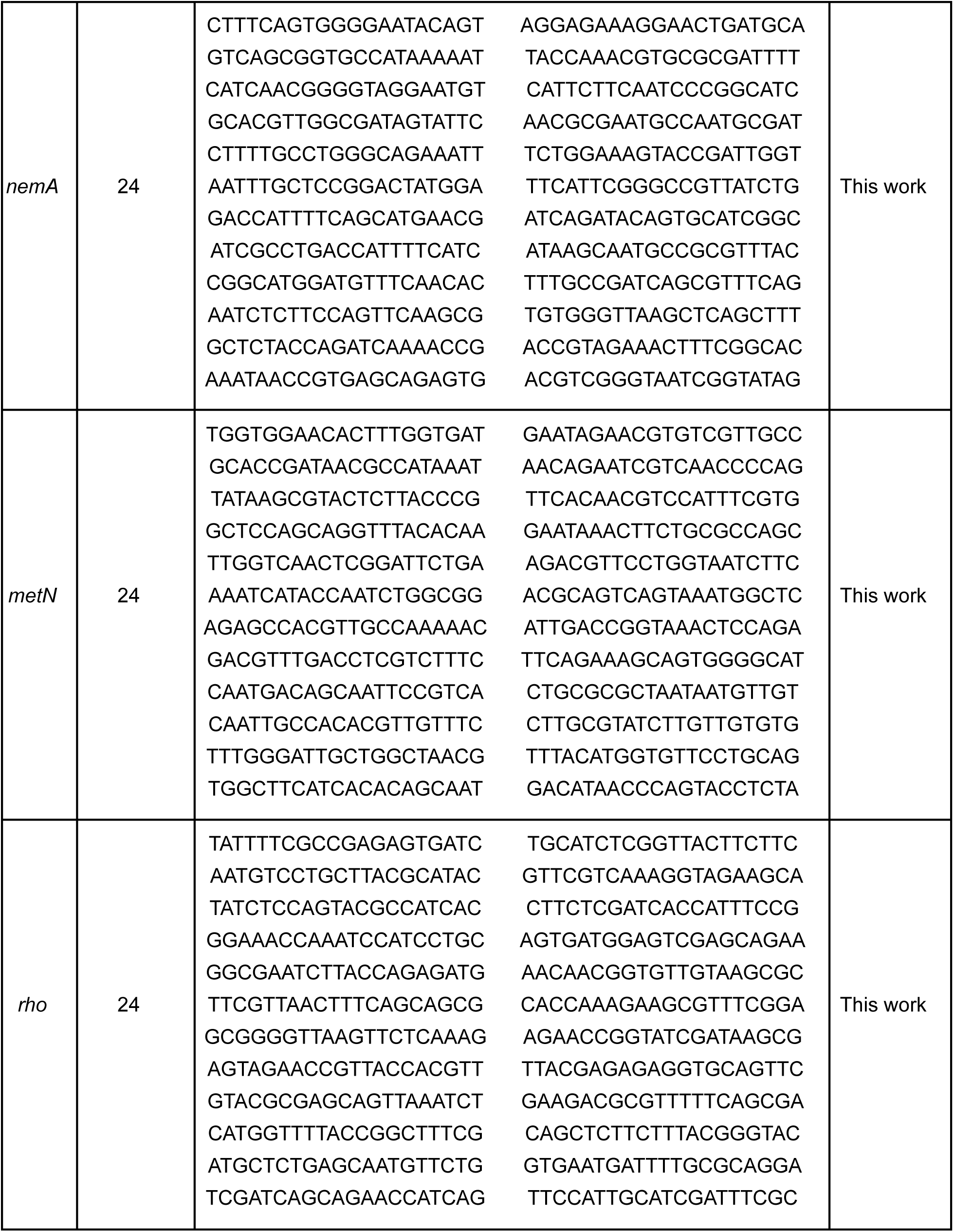
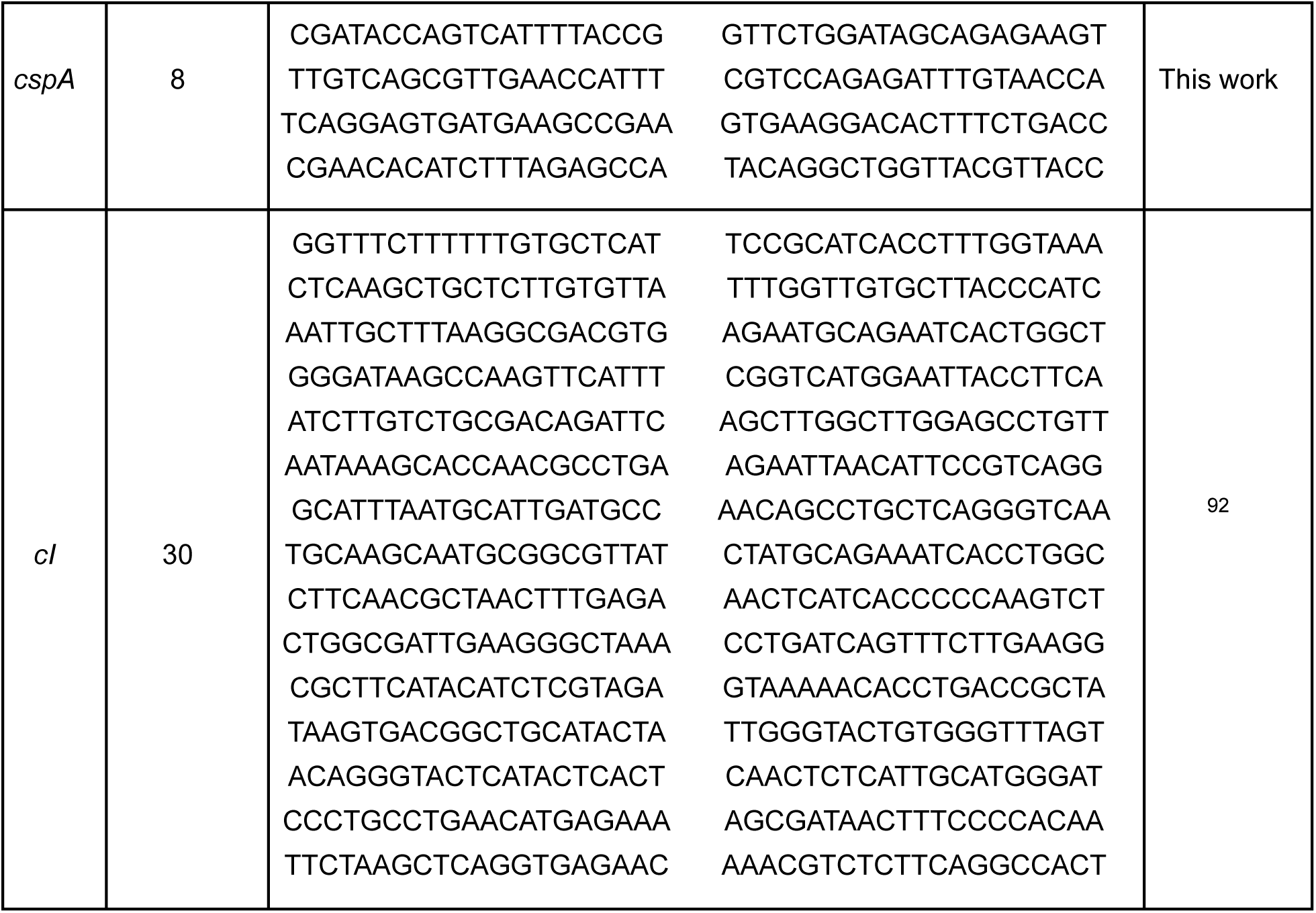
DNA oligos used for smFISH.

**Table S3:** Sample sizes for smFISH datasets.

| Gene | Number of cells in smFISH dataset 1 | Number of cells in smFISH dataset 2 |
| --- | --- | --- |
| *dnaA* | 2701 | 1772 |
| *nrdA* | 1481 | 1203 |
| *nemA* | 1077 | 2582 |
| *metN* | 1370 | 1892 |
| *rho* | 2113 | 823 |
| *cspA* | 572 | 1772 |
| *cl* (Negative control) | 841 | 1309 |

**Table S4:** Evidence of repressed state in high-amplitude cell cycle expression clusters. Evidence that genes within E. coli clusters Ec9 and Ec17 (Fig. 5C & D) are autorepressed or otherwise in a repressed state. Besides the sources listed, the EcoCyc^74,93^ database was used as a major source of information.

| Gene ID | Gene name | Description | Cluster | Operon | Regulation/evidence of repression | Ref. |
| --- | --- | --- | --- | --- | --- | --- |
| b3872 | *yihL* | Putative DNA-binding transcriptional regulator | Ec17 | *yihLM* | Nac-repressed; *yihL* is a GntR-family regulator so may have repressor function; *yihM* is induced by hexane so may have specific regulation | 94,95 |
| b4017 | *arpA* | Regulator of acetyl CoA synthetase | Ec17 | *arpA* | Unknown, but gene immediately downstream of the autorepressed transcription factor *iclR* |  |
| b4018 | *iclR* | DNA-binding transcriptional repressor IclR | Ec17 | *iclR* | Autorepression (also represses *aceBAK* operon) | 96 |
| b4191 | *ulaR* | DNA-binding transcriptional repressor UlaR | Ec17 | *ulaR* | Regulation unknown but repressor of *ulaG* and *ulaBCDEF* operons | 97 |
| b4278 | *insG* | KpLE2 phage-like element; IS4 putative transposase | Ec17 | *insG* | Unknown but NanR repressor binds promoter | 98 |
| b1650 | *nemA* | N-ethylmaleimide reductase | Ec17 | *nemRA-gloA* | Operon autorepressed by NemR (*gloA* partially transcribed by read-through from this operon) | 31–33 |
| b3502 | *arsB* | Arsenite/antimonite:H<sup>+</sup> antiporter | Ec17 | *arsRBC* | Operon autorepressed by ArsR | 99 |
| b4014 | *aceB* | Malate synthase A | Ec9 | *aceBAK* | Repressed by IclR; repressed by CRP in the presence of glucose | 100 |
| b4015 | *aceA* | Isocitrate lyase | Ec9 | *aceBAK* | Repressed by IclR; repressed by CRP in the presence of glucose | 100 |
| b4016 | *aceK* | Isocitrate dehydrogenase kinase/isocitrate dehydrogenase phosphatase | Ec9 | *aceBAK* | Repressed by IclR; repressed by CRP in the presence of glucose | 100 |
| b2675 | *nrdE* | Ribonucleoside-di phosphate reductase 2, $\alpha$ subunit dimer | Ec9 | *nrdHIEF* | Repressed by NrdR; repressed by FUR in the presence of iron | 101,102 |
| b2676 | *nrdF* | Ribonucleoside-di phosphate reductase 2, $\beta$ subunit dimer | Ec9 | *nrdHIEF* | Repressed by NrdR; repressed by FUR in the presence of iron | 101,102 |
| b3574 | *plaR* | DNA-binding transcriptional repressor PlaR | Ec9 | *plaR* | Autorepression (also represses L-lyxose catabolism operon) | 103 |
| b3605 | *lldD* | L-lactate dehydrogenase | Ec9 | *lldPRD* | Autorepression by LldR within the same operon | 104 |

